# Lateral Propagation from Sensory Cortex Provides Differential Gain for Sensory Selection

**DOI:** 10.64898/2026.10.23.761374

**Authors:** Angelina Lam, Sarah Kabbara, Alexandra Tejada, Hongdian Yang, Erfan Nozari, Edward Zagha

## Abstract

Selectively responding to target stimuli while ignoring distractors is a fundamental cognitive ability. In neocortex, such selection is associated with enhanced responses to target stimuli relative to distractor stimuli, yet the gain mechanisms underlying this differential responsivity are poorly understood. Most proposed gain mechanisms emphasize modulation within the center response field of a stimulus. In contrast, here we investigate modulation of lateral propagation from the center response field as a potential gain mechanism during a whisker-based selective detection task in mice. Both target (Go) and distractor (NoGo) stimuli evoked activity that propagated beyond their center response fields, with substantially greater lateral propagation for target stimuli. Target-evoked propagation increased with task learning and, in expert mice, exhibited the spatiotemporal characteristics of a traveling cortical wave preferentially involving infragranular layers. Furthermore, we demonstrate that frontal cortex actively suppresses distractor-evoked sensory responses, including both center region activity and lateral cortical recruitment. Computational modeling showed that modulation of lateral excitation could reproduce key features of these experimental findings. Together, our results support the regulation of lateral cortical propagation as a mechanism for differentially amplifying behaviorally relevant sensory representations during sensory selection.

## Introduction

A hallmark of goal-directed behavior is the ability to respond to task-relevant stimuli while ignoring distracting stimuli, a cognitive construct known as sensory selection. A longstanding framework for sensory selection proposes that distractor representations are attenuated while behaviorally relevant stimuli are preferentially transmitted for further processing (Treisman, 1964). Neural recordings during task performance have localized such attenuation within neocortex across diverse tasks and species (Aruljothi et al., 2020; Mante et al., 2013; Moran and Desimone, 1985; Sridharan et al., 2014; Wiederman and O’Carroll, 2013). A common observation is that differences between target-evoked and distractor-evoked responses become progressively greater from lower-order to higher-order cortical areas.

Less well established are the neural mechanisms that generate this differential responsivity. A dominant hypothesis is that sensory selection depends on differential gain modulation that amplifies target stimuli and/or suppresses distractor stimuli across sequential processing stages. However, the form of this gain modulation remains heavily debated (Fig. 1D-G). One possible mechanism is gating within thalamus along ascending sensory pathways before signals reach neocortex (Crick, 1984; McAlonan et al., 2008; Rodenkirch et al., 2019; Wang et al., 2010; Wimmer et al., 2015). Another mechanism is top-down amplification and/or suppression of sensory responses within neocortex (Carrasco, 2011; Maunsell and Treue, 2006; Maunsell, 2015; Reynolds and Chelazzi, 2004; Reynolds et al., 2000; Treue and Trujillo, 1999). A third proposes changes in spike synchronization across populations of cortical neurons, thereby altering their impact on downstream regions (Fries, 2009; Fries et al., 2001; Fries et al., 2002).

**Figure 1.**
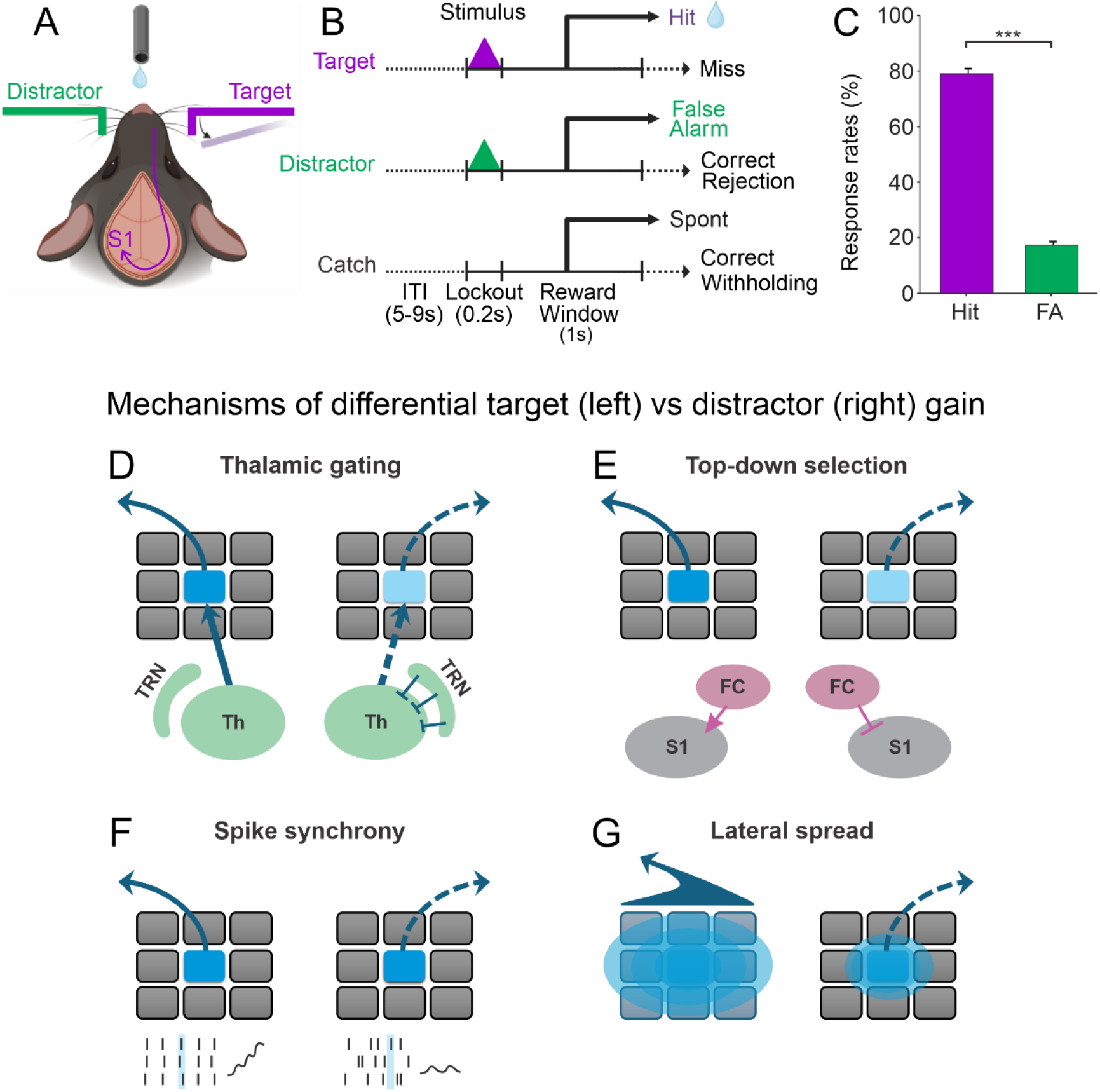
Potential mechanisms of differential target versus distractor gain during sensory selection. (A) Schematic of the whisker-based Go/NoGo selective detection task. Target and distractor stimuli are brief deflections of opposite whisker fields. (B) Trial structure and behavioral contingencies. Licking during the response window on target trials is scored as a hit and rewarded with a water reward; licking on distractor trials is scored as a false alarm and punished with a timeout. (C) Hit and false-alarm rates during expert performance. (D-G) Candidate mechanisms for differential target versus distractor gain, with each rectangular region reflecting a cortical column. (D) Thalamic gating of ascending sensory input (Th: Thalamus, TRN: Thalamic Reticular Nucleus), (E) top-down modulation of activity within the center response field (FC: Frontal Cortex, S1: Primary Somatosensory Cortex), (F) modulation of local spike synchrony, and (G) modulation of lateral propagation from the center response field into surrounding cortex. Data in C are mean +/- SEM. ***, p=6.87x10^-44^.

Canonical formulations of these mechanisms focus on activities within the center response field of a stimulus (i.e., for neurons specifically tuned to the presented stimulus). And yet sensory responses in neocortex are not confined to their center response fields. This phenomenon is particularly well established in whisker primary somatosensory cortex (wS1) in rodents. Whereas neurons in thalamorecipient layer 4 respond preferentially to their principal whisker, neurons in other cortical layers exhibit substantial multi-whisker responses, reflecting lateral recruitment beyond the initial cortical representation (Armstrong-James et al., 1992; Brett-Green et al., 2001; Chen-Bee et al., 2012; Clancy et al., 2015; Ferezou et al., 2007; Frostig et al., 2008; Hooks et al., 2011; Lefort et al., 2009; Masino and Frostig, 1996; Masino et al., 1993; Petersen, 2007; Simons, 1978). Foundational imaging studies further demonstrated that this lateral propagation is highly context dependent, varying with behavioral state and stimulus conditions, including anesthesia versus wakefulness and passive versus active touch (Ferezou et al., 2006; Lee et al., 2020; Peron et al., 2015). Selective regulation of this lateral recruitment could therefore provide a distinct form of cortical gain, amplifying the population representation of behaviorally relevant stimuli beyond their center response fields. Here, we investigate this possibility by examining how lateral propagation changes with task learning and how frontal cortex regulates target-evoked and distractor-evoked propagation.

To investigate these hypotheses, we used a Go/NoGo sensory selection task in mice (Aruljothi *et al*., 2020; Marrero et al., 2024; Marrero et al., 2021; Zareian et al., 2023; Zareian et al., 2021; Zhang and Zagha, 2023). In this task, Go (target) and NoGo (distractor) stimuli are presented on separate trials as rapid whisker deflections in opposite whisker fields (Fig. 1A-C). As in classic studies of sensory selection, we previously observed progressive filtering of target and distractor representations across neocortex: neuronal responses to the two stimuli were similar in wS1, yet only target stimuli effectively recruited frontal cortex (Aruljothi *et al*., 2020). Here, we find that lateral propagation from wS1 is differentially regulated during sensory selection. Target-evoked propagation increases with learning and, in expert animals, develops features of a traveling cortical wave in infragranular layers, whereas frontal cortex actively suppresses distractor-evoked sensory responses, including their lateral cortical recruitment. Computational modeling further shows that modulation of lateral excitation can reproduce key features of these experimental findings. Together, these results support regulation of lateral cortical propagation as a mechanism of differential sensory gain during sensory selection.

## Results

In our Go/NoGo selective detection task, mice are presented with target trials or distractor trials, consisting of rapid whisker deflections on opposite sides of the snout (Fig. 1A). If mice respond (lick) to target trials, they receive a water reward (Hit); if they respond to distractor trials, they are punished with a timeout (False Alarm) (Fig. 1B). We consider mice expert in this task once they achieve a hit rate vs false alarm rate d’>1 for three consecutive days, which typically occurs within two weeks of daily training. The expert sessions analyzed in this study (n=76 behavioral-recording sessions from n=21 mice) had the following behavioral profile: hit rate: 79 +/- 2%, false alarm rate: 17 +/- 1%, d’: 2.0 +/- 0.1 (paired t-test comparing hit rate and false alarm rate: t(75) = 30.37, p = 6.87x10^-44^; Fig. 1C).

### Target and distractor stimuli evoke similar initial sensory encoding within their center response fields

We focus all analyses (here and throughout) on the lockout period, from the onset of the stimulus to the onset of the response window, and exclude any trials with licking responses during the lockout. We previously reported single unit spiking activities of wS1 neurons aligned to the target stimulus on target trials (target_wS1) or aligned to the distractor stimulus on distractor trials (distractor_wS1) (Zhang and Zagha, 2023). Here, we highlight key findings from that dataset that provide insights into the neural mechanisms of sensory gain. Target_wS1 and distractor_wS1 populations showed similar rising phases and peak single unit stimulus encoding (n = 415 and 230 units, respectively; Supplemental Fig. 1B). These data argue against differential target vs distractor filtering along the sub-cortical ascending pathway (Fig. 1D) or early top-down selection within wS1 (Fig. 1E). However, there were differences suggestive of differential wS1 stimulus processing. Compared to distractor_wS1 neurons, target_wS1 neurons displayed increased pre-stimulus activity and sustained post-stimulus activity (Supplemental Fig. 1A).

Next, we performed new (unpublished) spike timing analyses to determine if population synchrony was selectively enhanced for target_wS1 compared to distractor_wS1 populations (n=17 target_wS1 recording sessions and n=10 distractor_wS1 recording sessions; Supplemental Fig. 2). We used Spike Time Tiling Coefficient (STTC) analyses, which quantify pair-wise spike synchrony from simultaneously recorded units while minimizing confounds of spike rate (see Methods). Results shown in Supplemental Fig. 2 and described below used a 15 ms synchrony window, and similar findings were obtained from 5 ms and 25 ms synchrony windows (data not shown). For both target_wS1 and distractor_wS1 populations, spike synchrony was significantly above chance levels throughout the prestimulus and post-stimulus epochs (Supplemental Fig. 2B,C and quantified for the pre-stimulus epoch in Supplemental Fig. 2D). Spike synchrony distributions were not significantly different either pre-stimulus (target_wS1, mean 0.073 +/- 0.007; distractor_wS1, mean 0.072 +/- 0.009; unpaired t-test: t(25) = 0.018, p = 0.99; Supplemental Fig. 2E) or peri-stimulus (target_wS1, mean 0.106 +/- 0.013; distractor_wS1, mean 0.125 +/- 0.024; unpaired t-test: t(25) = -0.782, p = 0.44; Supplemental Fig. 2F). Interestingly, spike synchrony in the target_wS1 population dipped below pre-stimulus levels after the target stimulus (Supplemental Fig. 2B). Consequently, spike synchrony was significantly *lower* in the target_wS1 post-stimulus condition compared to the distractor_wS1 post-stimulus condition (target_wS1, mean 0.049 +/- 0.005; distractor_wS1, mean 0.092 +/- 0.014; unpaired t-test: t(25) = -3.584, p = 0.0014; Supplemental Fig. 2G). Overall, these analyses indicate that spike synchrony modulation within wS1 (Fig. 1F) is unlikely to account for differential target vs distractor selection.

### Lateral propagation from sensory cortex is enhanced for target compared with distractor stimuli

In GCaMP-expressing mice, we paired expert task performance with widefield Ca^2+^ imaging of dorsal neocortex. We previously analyzed the propagation of target and distractor responses to frontal cortex (Aruljothi *et al*., 2020). Here, we instead focus on propagation during the lockout period within parieto-occipital cortex. We highlight an example expert behavioral-recording session (Fig. 2A,B). Within this session there were similar peak activations within the center response fields: target_wS1 region on target trials and distractor_wS1 region on distractor trials (neurometric d’: target_wS1, 0.96; distractor_wS1, 0.98). For distractor stimuli (Fig. 2B), the activity remained largely confined to wS1, decaying to baseline (d’=0) within approximately 1 mm from the center response field. In contrast, for target stimuli (Fig. 2A), the activity propagated broadly throughout parieto-occipital cortex bilaterally, in addition to activating frontal cortices.

**Figure 2.**
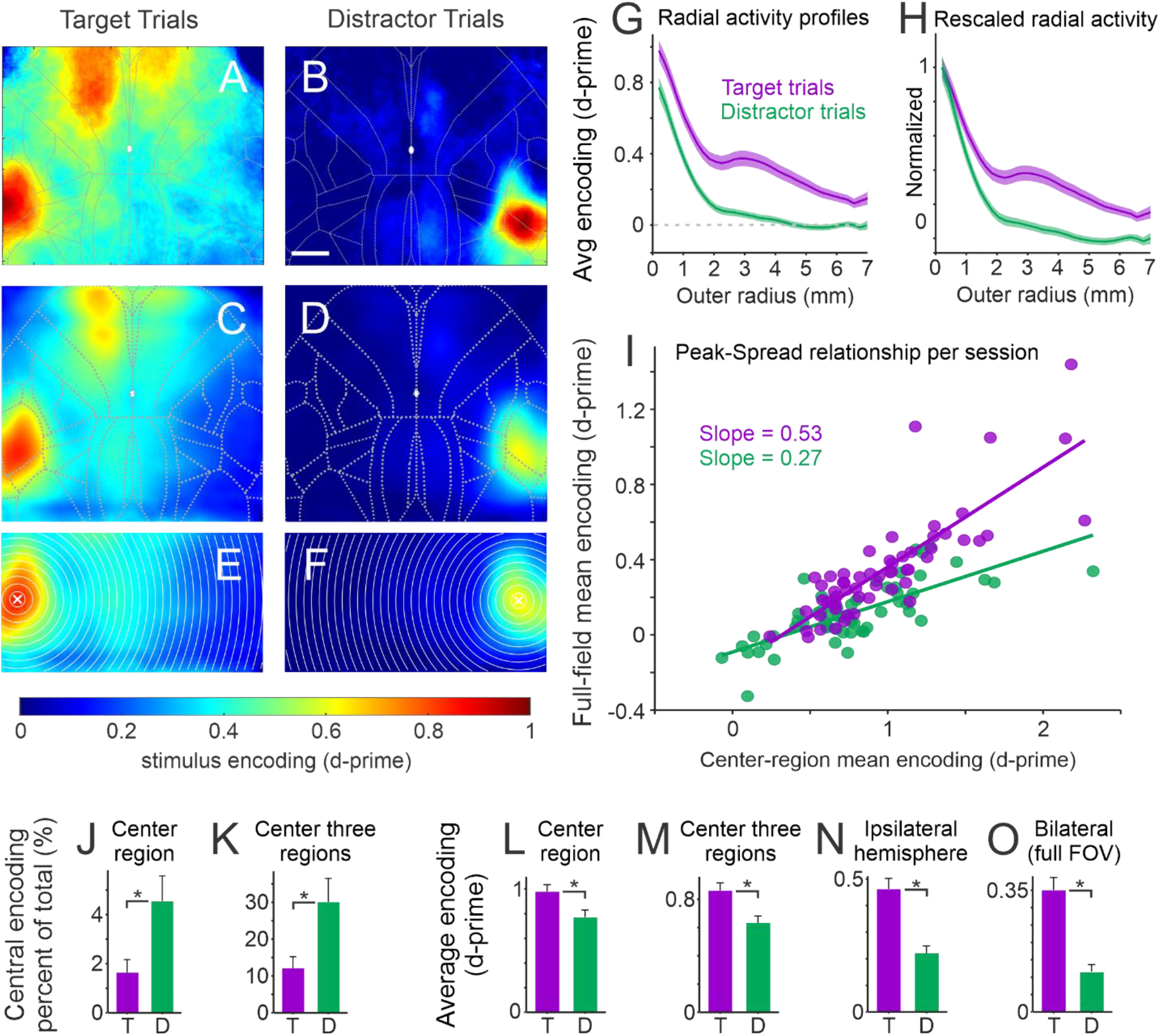
Lateral propagation from sensory cortex is enhanced for target stimuli compared to distractor stimuli. (A,B) Example widefield Ca²⁺ imaging session showing stimulus encoding (neurometric d′) after target (A) or distractor (B) stimulus onset. Target and distractor stimuli evoke similar peak activation within their respective wS1 center response fields, whereas target-evoked activity propagates more broadly across parieto-occipital cortex. (C,D) Grand-average stimulus encoding maps across expert recording sessions. (E,F) Radial-ring analysis centered on the location of maximum stimulus encoding for target (E) and distractor (F) trials. (G) Mean radial activity profiles showing greater peak activation and spatial propagation of target-evoked activity. (H) Radial profiles normalized to center-region activity, demonstrating more gradual spatial decay for target responses. (I) Relationship between center-region (peak) and full-field (spread) mean encoding across sessions. The steeper target trial slope indicates greater full-field recruitment for a given center response. Note, ‘full-field’ refers to the parieto-occipital region depicted in (E,F) and excludes frontal cortex. (J,K) Fraction of total parieto-occipital activation contained within the center response field (J) or center three radial regions (K). (L-O) Mean stimulus encoding measured within progressively larger spatial regions: center response field (L), center three regions (M), ipsilateral hemisphere (N), and bilateral parieto-occipital field (O). Lines/shading and bars denote mean +/- SEM. T, target; D, distractor. *, p<0.05. Scale bar in (B), 1 mm.

Fig. 2C,D show grand averages from all expert sessions (n=58 sessions). To quantify the spatial propagation of cortical activity, we conducted a radial ring analysis on parieto-occipital cortex of the neurometric d’ maps (Fig. 2E,F). For each session, we first identified the region with maximum activity, which we distinguished as the center response field. This center region has a radius of 205 µm which contains approximately 2 to 3 barrel column areas, likely reflecting the somatotopic region of the deflected whiskers. Radial activity profiles were generated for each session and subsequently averaged across sessions (Fig. 2G). As evident in the normalized profiles (Fig. 2H), the spatial decay of stimulus encoding on target trials was more gradual than on distractor trials.

To capture the peak activity vs lateral spread relationship, we plotted the peak encoding of the center region against the full-field mean encoding across parieto-occipital cortex per session (Fig. 2I). Target trial and distractor trial data were each well fit by linear regression (target trials: R^2^ = 0.68, p = 2.3x10^-15^; distractor trials: R^2^ = 0.56, p = 1.7x10^-11^), reflecting a strong correlation between peak activation and total activation. However, the slope of the peak-spread relationship was greater for target trials compared to distractor trials (target trials: slope = 0.53; distractor trials: slope = 0.27; mixed-effects regression, center activity × trial type interaction: F(1,112)= 19.0, p = 3.0x10^-5^; Fig. 2I). To gain insight into these findings, we conducted numerical simulations (Supplemental Fig. 3). We simulated spatial activation as a Gaussian distribution. Multiplicative scaling of the Gaussian distribution did not change the peak-spread relationship (Supplemental Fig. 3A,B). In contrast, increasing the standard deviation of the distribution increased the slope of the peak-spread relationship (Supplemental Fig. 3C,D), analogous to the target trial data (Fig. 2I). Thus, we conclude that the increased lateral propagation on target trials is not simply a consequence of stronger activation at the center response field but instead reflects a difference in propagation from the center into the surrounding regions.

We conducted a series of quantifications to better characterize the spatial activation. First, we determined the fraction of total parieto-occipital activation accounted for by the center response field. Remarkably, the center response field accounted for only 1.6 +/- 0.5 % of the total activation on target trials and 4.5 +/- 1.0 % of the total activation on distractor trials (paired t-test, t(57) = 2.61, p=0.012, n=58 sessions; Fig. 2J). Fractional activation of the center three rings, corresponding to approximately 20 barrel columns which is well beyond the somatotopic region of direct stimulation, still accounted for a minority of the total activation (target trials, 12.1 +/- 3.1 %; distractor trials, 30.1 +/- 6.4 %; paired t-test, t(57) = 2.63, p=0.011). We also compared the magnitude of activation between target and distractor trials. At the center response field, activation was on average 21% lower for distractor compared to target trials (paired t-test, t(57) = -3.11, p=0.0029; Fig. 2L). Differences in neural activation increased with increasing window size (Fig. 2M-O), and was 67% lower for distractor trials when considering the entire parieto-occipital field of view (paired t-test, t(57) = -5.82, p=2.9x10^-7^; Fig. 2O).

### Enhanced target-evoked lateral propagation emerges with task learning

To determine the emergence of differential lateral propagation, we recorded neural activity longitudinally across task learning (Fig. 3) (Marrero *et al*., 2024). Recordings were conducted across naïve (discrimination d’<0.8), intermediate (0.8 < d’ < 1.2), and expert (d’ > 1.2) sessions (n=6 mice). For target trials, peak encoding within the center region remained remarkably consistent across training (Fig. 3A-D). From intermediate to expert performance, however, there was a notable increase in the lateral propagation from the center response field (Fig. 3B-E). Accordingly, there was a significant increase in the slope of the peak-spread relationship across learning stages (mixed-effects regression, center activity × learning stage: F(2,12) = 8.30, p = 0.0055; Fig. 3F).

**Figure 3.**
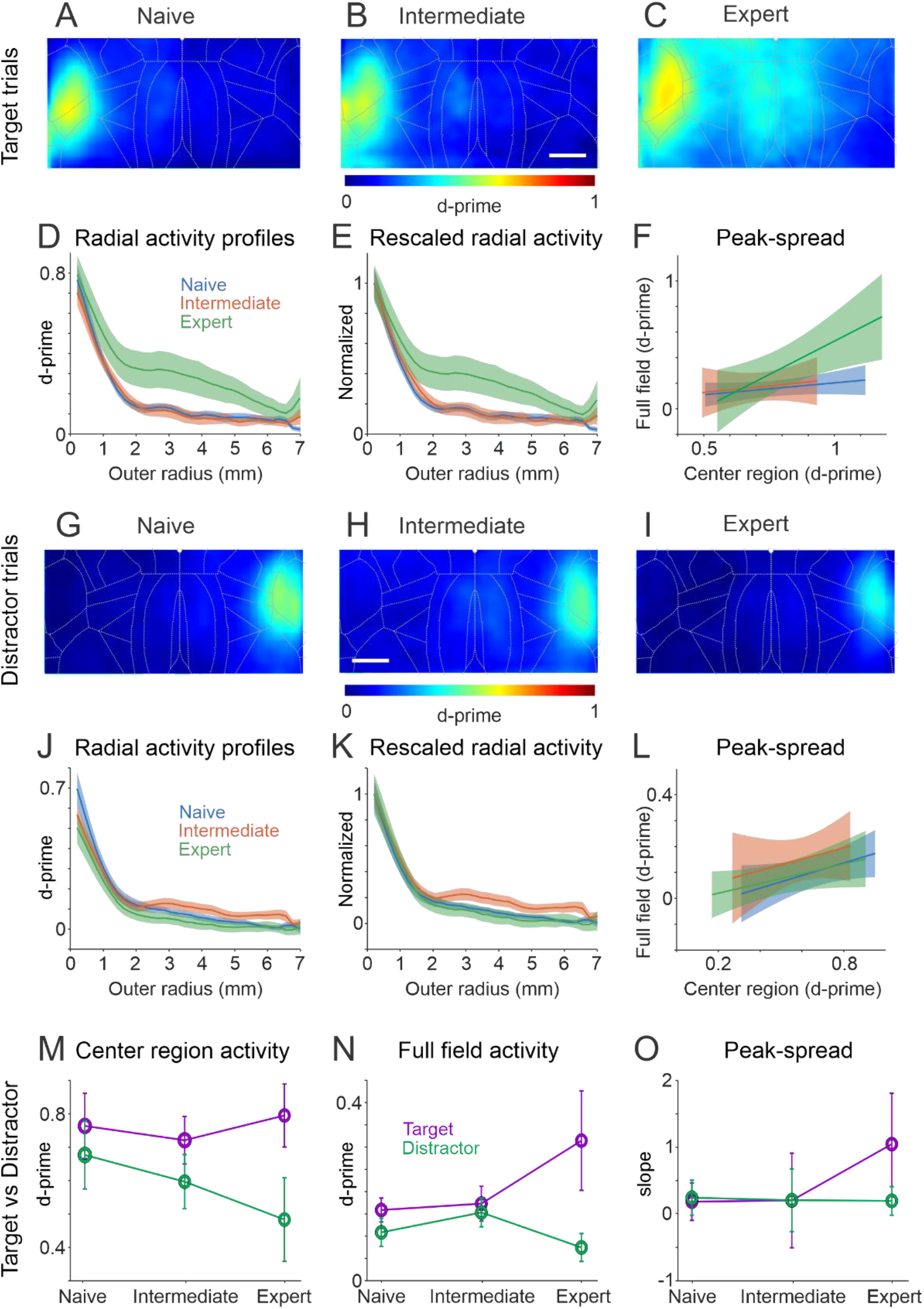
Enhanced target-evoked lateral propagation emerges with task learning. (A-C) Grand-average target-evoked stimulus-encoding maps from parieto-occipital cortex during naïve (A), intermediate (B), and expert (C) performance. Note the robust increase in lateral propagation in expert sessions compared to modest changes in peak encoding within the center field. (D) Radial target-evoked activity profiles across learning stages. (E) Radial profiles normalized to center-region activity. (F) Peak-spread relationships across learning stages, showing a steeper relationship for expert sessions. (G-L) As in (A-F) but for distractor trials, demonstrating overall reductions in activity across training. (M-O) Comparison of target and distractor responses across training stages for the center region (M), full-field (N), and the slope of the peak-spread relationship (O). Radial-profile shading and error bars denote SEM; regression shading denotes the 95% confidence interval of the fitted mean. Scale bars in (B, H), 1 mm.

Different learning-related dynamics were observed for distractor trials. There was a trend towards decreasing activity within the center region with training (Fig. 3G-I), as well as a trend towards a transient increase in lateral propagation in intermediate sessions (Fig. 3G-K). However, unlike for target trials, for distractor trials there was not a significant change in the slope of the peak-spread relationship across learning stages (mixed-effects regression, center activity × learning stage: F(2,12) = 0.85, p=0.45; Fig. 3L).

We further compared these metrics between target and distractor trials across training stage (Fig. 3M-O). For center region and full-field activity, trial type × training stage interactions did not reach significance (2 × 3 repeated-measures ANOVA, Greenhouse–Geisser corrected; center region: F(1.16,5.81) = 2.30, p = 0.18; full field: F(1.05,5.26) = 3.40, p = 0.12). In contrast, the peak-spread relationship showed a significant trial type × training stage × center activity interaction (mixed-effects regression, F(2,24) = 14.32, p = 8.1x10^-5^; Fig. 3O). These analyses indicate that lateral propagation increases for target trials late in training, as mice acquire expert performance. In contrast, lateral propagation does not increase for distractor trials, which may be due to lack of selection and/or active suppression, which we address in Fig. 6.

### Target-evoked activity propagates from wS1 with the spatiotemporal characteristics of a cortical traveling wave

Limitations of widefield Ca^2+^ imaging include slow timescales and restricted depth resolution. Therefore, to further resolve the spatio-temporal properties of lateral propagation on target trials we used laminar multi-electrode array recordings (Fig. 4). Within the wS1 center response field (Fig. 4A), we observed short-latency, negative-going local field potentials (LFPs) with the largest amplitudes in intermediate layers, as expected. As we extended the recordings medially (Fig. 4B-E), we observed shifts in stimulus-evoked LFP latency, polarity, and depth profile. LFP signals in deeper layers became more prominent, and the LFP polarity transitioned from negative-going to positive-going. Current source density (CSD) analyses of wS1 recordings revealed early current sinks in thalamorecipient layer IV (0.4-0.6 mm deep, Fig. 4F,G). Interestingly, at 1.5 mm medial, we observed robust current sinks localized to infragranular layers (Fig. 4H,I and Supplemental Fig. 4).

**Figure 4.**
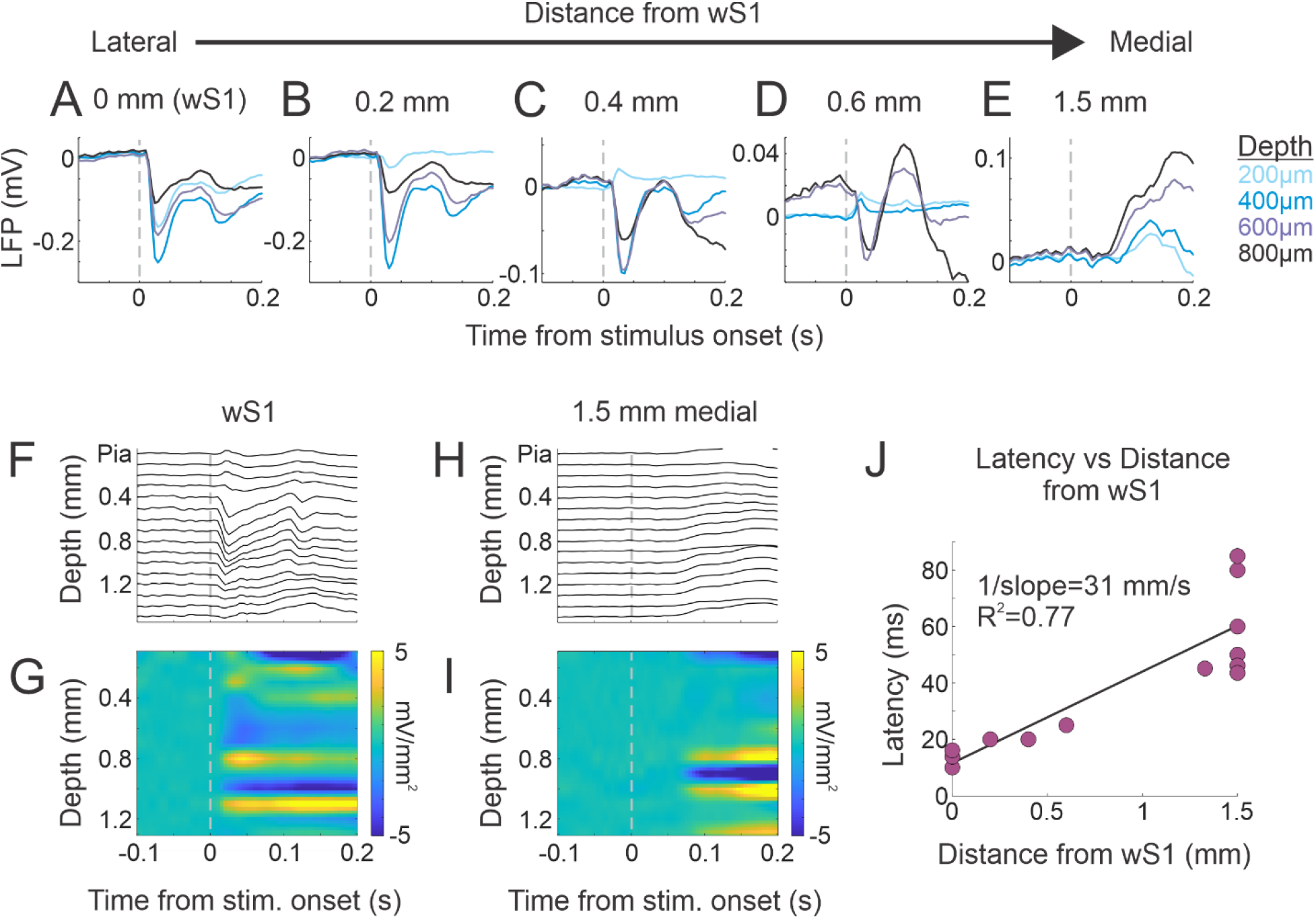
Target-evoked activity propagates medially from wS1 with the spatiotemporal characteristics of a cortical traveling wave. (A-E) Target-evoked LFP responses recorded at increasing medial distances from the wS1 center response field. Traces show responses across recording depths. With increasing distance from wS1, LFP response latency increased and the laminar distribution and polarity of the response changed. (F,G) Laminar LFP (F) and current-source-density (CSD) (G) responses from an example wS1 session, showing an early sink in thalamorecipient layer IV (∼0.4-0.6 mm deep). (H,I) Laminar LFP (H) and CSD (I) responses from an example 1.5 mm medial session, where the prominent stimulus-evoked sink is localized to infragranular layers. (J) LFP response latency as a function of distance from wS1. Linear regression yielded R²=0.77 and an estimated propagation velocity of 31 mm/s (inverse of the slope). These observations are consistent with a target-evoked cortical wave propagating from the sensory cortex center response field, preferentially involving infragranular layers.

Latency to LFP response onset increased from 10-15 ms within wS1 to 45-85 ms at 1.5 mm medial. To estimate the speed of lateral propagation, we plotted the latency vs distance for each behavioral-recording session (Fig. 4J). Latency increased linearly with distance (slope = 32.3 ms/mm, R^2^ = 0.77, p = 0.0002, n = 12 recording sites). The inverse of the regression slope yielded an estimated propagation speed of 31 mm/s, consistent with published estimates of cortical wave propagation (Liang et al., 2023; Yang et al., 2015). Together, these data are consistent with the target stimulus initiating a cortical wave from the center response field within wS1 that propagates laterally, preferentially within infragranular layers.

We interpret the enhanced lateral propagation on target trials as a potential mechanism to increase the gain of the target stimulus to facilitate triggering the associated behavioral response. This is in contrast to distractor trials, for which reduced gain may help prevent response triggering. However, we recognize an alternative explanation, that enhanced lateral propagation on target trials may be a consequence of movement on Go trials, which are largely suppressed on NoGo trials (Zagha et al., 2022). To distinguish between these possibilities, we compared the temporal alignments of various task-related events (Fig. 5). We distinguish ‘stimulus-related’ processes as short latency events that align to the stimulus regardless of reaction time, and ‘response-related’ processes as longer latency events that align to the response regardless of reaction time (Lam et al., 2026). In Fig. 5A we plot the stimulus- and response-alignment of the whisker paddle motion energy (ME) as an obviously stimulus-related process. The movement of the paddle occurred at a short and fixed latency from the stimulus onset (0.010 +/- 0 s, within the first post-stimulus imaging frame). In contrast, the paddle ME had a variable latency when aligned to the response. Similar dynamics were observed from the wS1 LFP signal (Fig. 5B), with the initial downward deflection showing a short and fixed latency from the stimulus onset (0.013 +/- 0.003 s).

**Figure 5.**
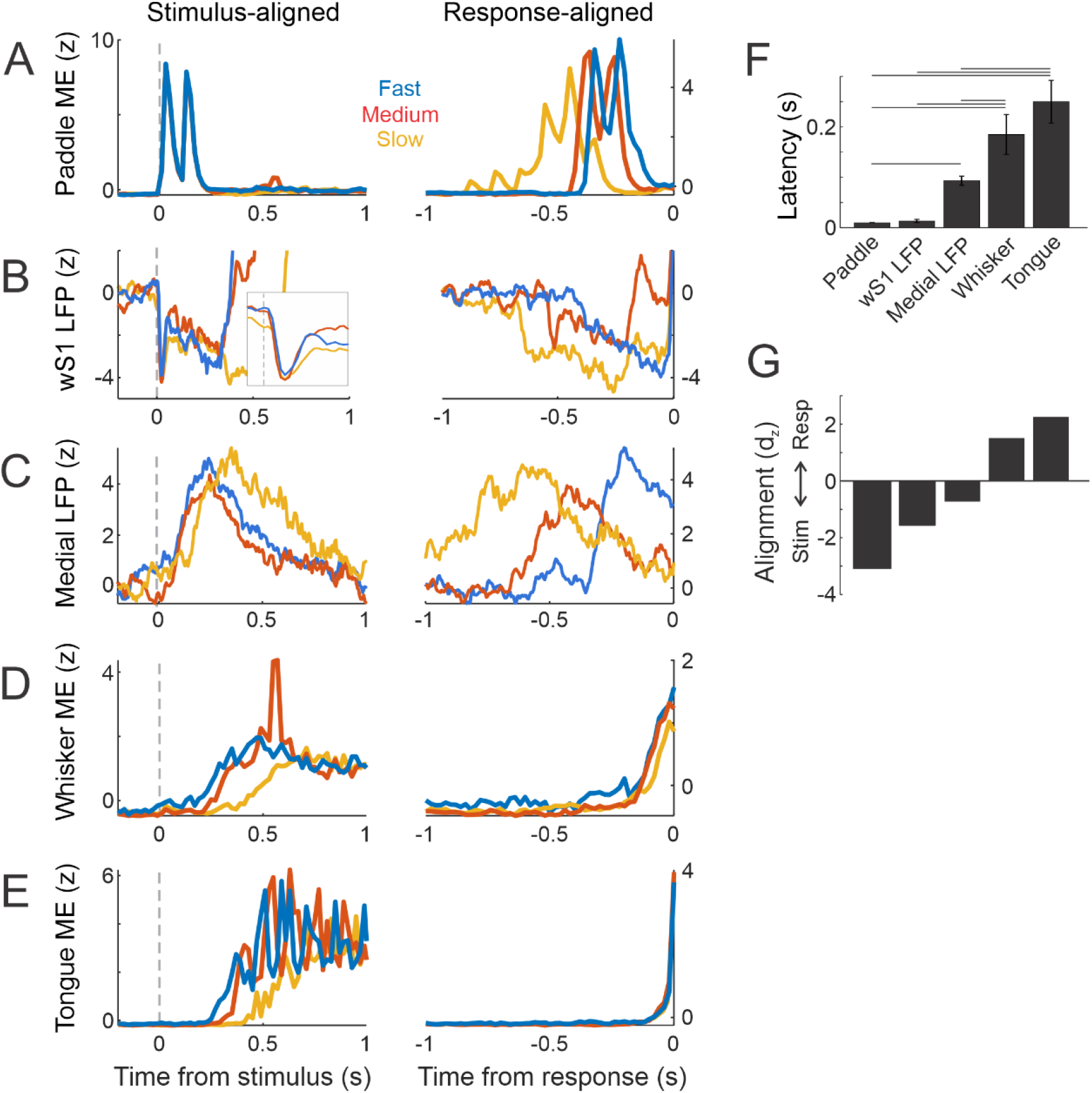
Cortical propagation is temporally coupled more strongly to sensory stimulation than to the behavioral response. (A-E) Example session data, with activity separated into fast, medium, and slow reaction time trials and aligned either to stimulus onset (left) or behavioral response (right). (A) Motion energy (ME) of the stimulus paddle provides a stimulus-locked reference signal. (B) wS1 LFP responses are similarly tightly stimulus aligned. (C) Medial parietal LFP responses occur later than wS1 responses but remain more consistently aligned to the stimulus than to the behavioral response. (D) Whisker and (E) tongue motion energy are progressively more tightly aligned to the behavioral response. (F-G) Session-average data. (F) Response latency across signals, illustrating the temporal progression from stimulus delivery through cortical activity to overt movement. Connecting lines indicate significant pairwise differences from post hoc comparisons. (G) Alignment index (dz) comparing variability of stimulus-aligned and response-aligned latencies. Negative values indicate stronger stimulus alignment and positive values indicate stronger response alignment. Together, these analyses place medial cortical propagation between initial wS1 activation and overt movements.

At the other end of the sensory-to-motor transformation is the tongue ME, included as an obvious response-related process. The tongue ME occurred at a long and variable latency to the stimulus onset (0.250 +/- 0.042 s), in contrast to the short and fixed latency preceding the response (Fig. 5E). Exploratory studies found that whisker movements (including those not directly stimulated by the paddle, Supplemental Fig. 5) precede licking and other body movements in our task (data not shown). Whisker ME also displayed greater temporal alignment to the response, consistent with a response-related process (Fig. 5D).

With these comparisons in place, we analyzed the temporal alignments of the medial LFP signals (Fig. 5C). Medial LFP signals demonstrated greater temporal alignment to the stimulus, consistent with a stimulus-related process, while not as fast or fixed as the paddle ME and wS1 LFP signals. Across the full datasets, response latency differed significantly across signal types (Fig. 5F), with post hoc comparisons supporting a temporal progression from paddle motion and wS1 activity through medial parietal activity to overt whisker and tongue movements. Consistent with this progression, medial LFP signals showed preferential stimulus alignment, whereas whisker and tongue movements showed preferential response alignment (Fig. 5G). Together, these analyses place medial parietal activation between initial wS1 activation and the earliest body movements, and are therefore inconsistent with medial cortical propagation being a consequence of overt movements on Go trials.

### Motor cortex suppresses distractor-evoked sensory cortical recruitment

Next, we sought to determine whether lateral propagation is subject to top-down modulation (Fig. 6). We previously showed that suppression of the whisker region of motor cortex in the distractor hemisphere (distractor_wMC) increases false alarms (Zareian *et al*., 2023; Zhang and Zagha, 2023) and enhances the propagation of distractor-evoked activity into target_wS1 (Zhang and Zagha, 2023). However, the effects of distractor_wMC suppression on the broader spatial profile of sensory cortical activity were unknown. Here, we combined muscimol suppression of distractor_wMC (Supplemental Fig. 6) with widefield Ca^2+^ imaging in expertly performing mice (n=6 mice).

**Figure 6.**
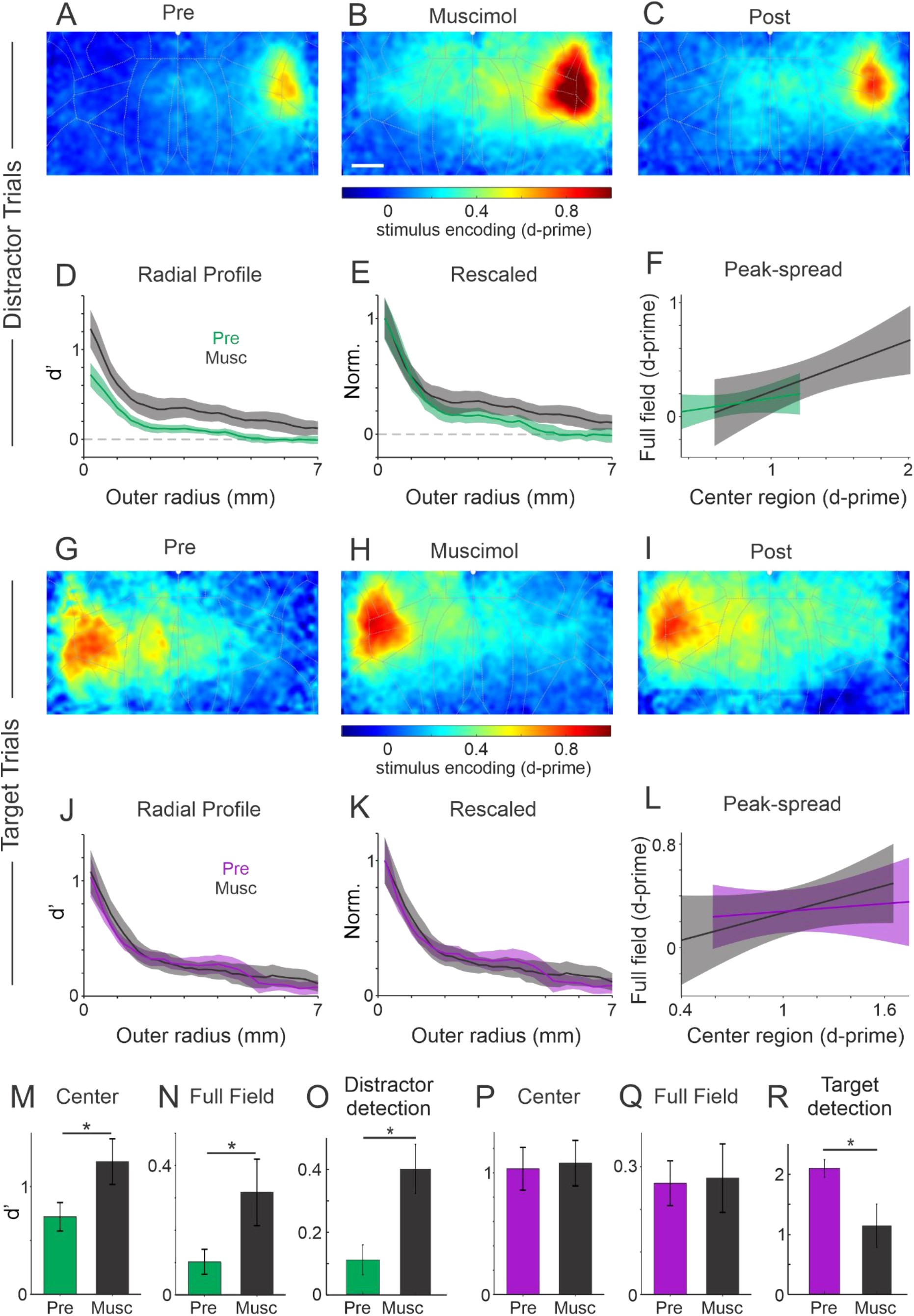
Suppression of distractor-aligned whisker motor cortex increases distractor-evoked sensory cortical recruitment. (A-C) Grand average distractor-evoked stimulus-encoding maps before (A), during (B), and after (C) muscimol suppression of distractor-aligned whisker motor cortex (distractor_wMC). distractor_wMC suppression increased distractor-evoked activity in wS1 and across surrounding parieto-occipital cortex, with partial recovery after muscimol washout. (D) Distractor-evoked radial activity profiles before and during distractor_wMC suppression. (E) Corresponding profiles normalized to center-region activity. (F) Peak-spread relationships before and during distractor_wMC suppression. (G-L) Corresponding analyses for target-evoked activity, which showed little change in spatial activation during distractor_wMC suppression. (M-R) Quantification of neural and behavioral measures before (color) and during (black) distractor_wMC suppression. (M,N) Mean distractor-evoked activity within the wS1 center region (M) and full-field (N). (O) Distractor stimulus detection. (P,Q) Mean target-evoked activity within the wS1 center region (P) and full-field (Q). (R) Target stimulus detection. Units in (M,N,P,Q) are neurometric d’; units in (O,R) are behavioral d’. Radial-profile shading and error bars denote SEM; regression shading denotes the 95% confidence interval of the fitted mean. *, p<0.05. Scale bar in (B), 1 mm.

distractor_wMC suppression substantially increased distractor-evoked activity in wS1 and across the surrounding parieto-occipital cortex (Fig. 6A,B), with recovery after muscimol washout (Fig. 6C). Radial-profile analyses similarly demonstrated increased activation across the parieto-occipital field (Fig. 6D,E). Consistent with disproportionate enhancement of broader cortical recruitment, distractor_wMC suppression significantly increased the slope of the peak-spread relationship (linear mixed-effects model, center activity × drug condition interaction: F(1,3.37) = 11.98, p = 0.034; Fig. 6F). In contrast, target trials recorded within the same sessions showed little change in spatial activation (Fig. 6G-K), and the peak-spread relationship was not significantly altered (center activity × drug condition interaction: F(1,8) = 1.33, p = 0.283; Fig. 6L).

distractor_wMC suppression increased distractor-evoked activity within the wS1 center region by 78% (paired t-test, t(5) = 3.34, p=0.021; Fig. 6M) and across the full field by 212% (paired t-test, t(5) = 2.58, p=0.0497; Fig. 6N). Behaviorally, distractor_wMC suppression increased distractor stimulus detection (Fig. 6O), consistent with the increased cortical representation of the distractor stimulus. In contrast, distractor_wMC suppression did not significantly alter target-evoked center region or full field activity (paired t-tests; center: t(5) = 0.41, p = 0.70; full field: t(5) = 0.14, p = 0.90; Fig. 6P,Q), despite reducing target stimulus detection (Fig. 6R). Together, these results demonstrate active top-down suppression of distractor-evoked sensory cortical activity, including recruitment beyond the wS1 center region, with disproportionate suppression of broader lateral cortical recruitment.

### Increased lateral coupling reproduces the spatial gain associated with target stimuli

Lastly, to examine potential mechanisms underlying the relationships between center response and lateral propagation, we simulated activity in a two-dimensional recurrent network (Fig. 7A). Each unit had recurrent interactions with itself (weight denoted as *W_s_*) along with lateral interactions with units within two lattice distances (*W_d_*_1_ and *W_d_*_2_; Fig. 7B). Network activity was driven by a uniform tonic background input and a focal ‘sensory’ input delivered to the center unit. We compared the baseline network with various modulations, in an attempt to qualitatively simulate the differences observed experimentally in distractor (baseline) and target (modulated) associated signaling. Specifically, for target compared to distractor trials, our experimental data support: enhanced center pre-stimulus baseline activity (Supplemental Fig. 1A), mildly enhanced center post-stimulus evoked activity (Fig. 2G,L), substantially enhanced full-field evoked activity (Fig. 2G,O), and enhanced spread-to-peak ratio (Fig. 2H,I).

**Figure 7.**
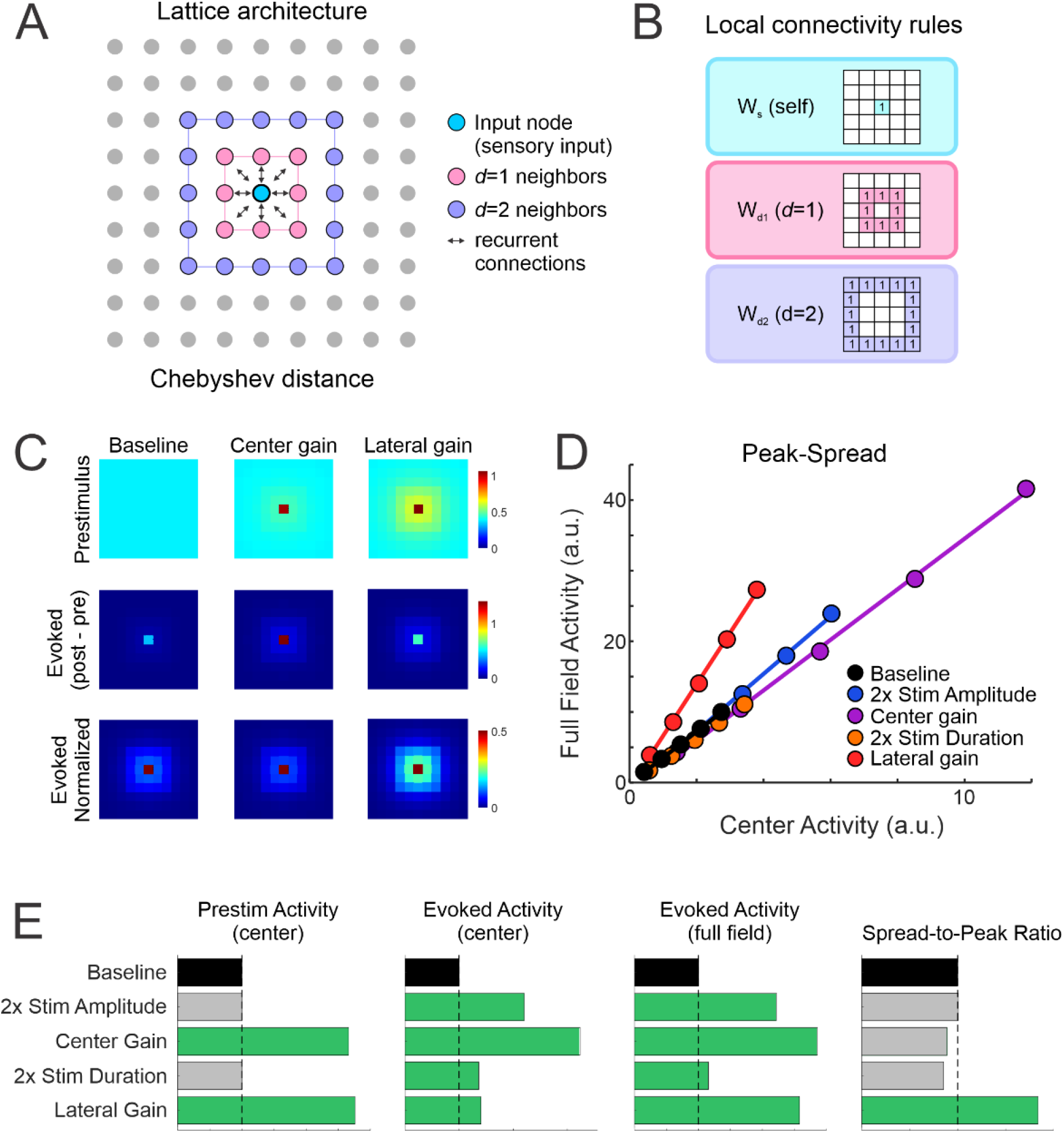
Increased lateral coupling reproduces the spatial gain associated with target stimuli. (A) Architecture of the two-dimensional recurrent network. A focal sensory input is delivered to the center unit; recurrent connections are defined according to Chebyshev distance. (B) Local connectivity weights, consisting of self-coupling (*W_s_*) and lateral coupling to units one (*W_d_*_1_) or two (*W_d_*_2_) lattice positions away. (C) Simulated prestimulus and stimulus-evoked activity for the baseline network and networks with increased center gain or lateral gain. Center gain was implemented by increasing self-coupling of the center unit, whereas lateral gain was implemented by increasing excitatory coupling between the center and surrounding units. (D) Center activity versus full field activity across five stimulus amplitudes for the baseline model and four gain manipulations. Increased lateral gain, but not the other model manipulations, produced a pronounced increase in the slope of the peak-spread relationship. (E) Summary across models of four experimentally observed target-associated features: prestimulus center activity, evoked center activity, evoked full-field activity, and spread-to-peak ratio. Green bars indicate changes in the experimentally observed direction. Only increased lateral gain qualitatively reproduces all four features of the target-associated condition.

In Fig. 7C, we simulate two potential gain mechanisms (see also Supplemental Fig. 7). ‘Center gain’ was simulated by increasing center unit self-coupling (*W_s_*), to simulate top-down enhancement of local excitability at the stimulated representation. ‘Lateral gain’ was simulated by increasing lateral-excitation to and from the center unit, to simulate enhanced lateral coupling between the stimulated representation and the surrounding cortices. Both center gain and lateral gain modulations increased prestimulus activity (Fig. 7C, top row). Increasing center gain enhanced stimulus-evoked center and surround activity (Fig. 7C, middle row) but largely preserved their relationship (Fig. 7C, bottom row). Notably, increasing lateral gain produced a disproportionate increase in stimulus-evoked surround activity (Fig. 7C, bottom row).

We included two additional modulations: increased stimulus amplitude to simulate pre-cortical gain (e.g., thalamic, Fig. 1D) and increased stimulus duration to simulate persistent target-evoked responses (Supplemental Fig. 1, (Sachidhanandam et al., 2013; Yang et al., 2016)). In Fig. 7D, we plot the peak-spread relationship for each model across 5 stimulus amplitudes. With the baseline condition as the comparison (black), *only the lateral gain modulation (red)* substantially increased the slope of the peak-spread relationship, as observed in our experimental data.

Across the five models we compare the four outcome measures identified in our experimental data (Fig. 7E, green indicating qualitative agreement with target compared to distractor associated outcomes). Notably, only the lateral gain modulation replicated all four outcomes. Together, these data support increases in lateral coupling as a potential mechanism of target-associated sensory gain.

## Discussion

In this study, we propose a mechanism of stimulus gain modulation based on the regulation of lateral cortical propagation during sensory selection. Mice were trained to respond to target stimuli while ignoring similar distractor stimuli presented on separate trials. Evoked responses to both target and distractor stimuli reached their corresponding somatotopic cortical regions, yet only target stimuli effectively propagated to frontal cortex (Aruljothi *et al*., 2020). Our neural recordings provide limited evidence that modulation within the center response field alone can account for target-distractor differences. In contrast, we observed robust differences in lateral propagation beyond the center response field. A hallmark of this enhanced propagation was a change in the peak-spread relationship (Fig. 2I), indicating disproportionately greater spatial recruitment as response magnitude increased. This relationship is inconsistent with purely uniform multiplicative gain and instead suggests preferential modulation of lateral cortical recruitment.

A prominent anatomic feature of neocortex is its recurrent intra-areal connectivity, including horizontal excitatory projections extending across neighboring cortical territories (Douglas and Martin, 2004). These connections provide a substrate through which focal sensory input can recruit populations beyond its center response field. Within the whisker system, lateral propagation has been shown to depend on both stimulus strength and behavioral state (Ferezou *et al*., 2007; Petersen et al., 2003). Recurrent cortical interactions have been implicated in various computations including contour integration, divisive normalization, and predictive processing (Carandini and Heeger, 2011; Field et al., 1993; Heeger, 1992; Muller et al., 2026; Stettler et al., 2002). Here, we propose an additional role for these interactions: regulating the spatial gain of sensory representations according to behavioral relevance.

One potential advantage of modulating lateral propagation is the ability to regulate the size and spatial extent of the recruited neuronal population. Center-field gain changes alter neurons already engaged by the stimulus and may saturate at high stimulus intensities (Carrasco, 2011; Fox et al., 2023; Maunsell and Treue, 2006; Maunsell, 2015; Reynolds and Chelazzi, 2004; Reynolds *et al*., 2000; Treue and Trujillo, 1999). In contrast, lateral propagation can recruit populations well beyond the center response field, which are not typically activated by the stimulus. In our task, deflection of only 2-3 target whiskers evoked activity across much of dorsal parietal cortex (Fig. 2A-H). Jacobs and Frostig (2017) noted that “the vast majority of whisker evoked activity in [wS1] occurs beyond columnar boundaries” (Jacobs and Frostig, 2017). Consistent with this observation, less than 2% of target-evoked widefield Ca^2+^ activity integrated across dorsal parieto-occipital cortex occurred within the center response field (Fig. 2J). Consequently, restricting analyses to center responses only risks overlooking a substantial component of sensory-evoked activity.

Previous studies have described wave-like propagation from wS1 following whisker stimulation (Ferezou *et al*., 2007; Gonzales et al., 2025; Jacobs and Frostig, 2017). Using high-resolution recordings, Gonzales et al. (2025) identified two temporally distinct whisker-evoked traveling waves within wS1. Based on its timing, the propagation observed here appears more similar to their “early” wave, although direct correspondence remains uncertain given the methodological and behavioral differences. Interestingly, wMC exerted distinct effects across the two studies: here, wMC suppressed distractor-evoked lateral propagation, whereas Gonzales et al. (2025) found that wMC facilitated the “late” traveling-wave component. These observations suggest that frontal input may regulate sensory propagation with temporal and behavioral specificity.

A notable feature of the wave-like responses observed here is their preferential expression within infragranular layers, based on CSD analyses. One candidate substrate is layer V pyramidal neurons, which possess extensive horizontal collateral axons (Staiger et al., 2016) and contribute to horizontal spread of recurrent cortical activity (Wester and Contreras, 2012). However, the present experiments do not establish that intracortical horizontal connections alone mediate the observed propagation. Additional pathways may contribute, including long-range recurrent loops through higher-order cortical areas (Manita et al., 2015; Minamisawa et al., 2018) and/or higher-order thalamus (Guo et al., 2020).

Increased lateral propagation on target trials emerged relatively late in training (Fig. 3), as mice stably expressed target-distractor discrimination. Whether similar modulation occurs during rapid attentional changes remains unknown, but previous studies provide suggestive parallels. Attending to a target stimulus alters receptive field properties of sensory cortical neurons (Connor et al., 1997; Fritz et al., 2003; Fritz et al., 2010; Moran and Desimone, 1985; Poort et al., 2015; Womelsdorf et al., 2006; Womelsdorf et al., 2008). Studies have demonstrated narrowing of target-aligned receptive fields and shifting of non-aligned receptive fields toward the target. These findings could reflect asymmetric changes in lateral propagation: increased propagation out of a target-aligned center response field for target stimuli and reduced propagation into a target-aligned center response field for distractor stimuli. Previous models have explained attentional receptive-field shifts as consequences of spatially structured modulation of sensory responses (Compte and Wang, 2006). These parallels suggest that the lateral propagation mechanism identified here during learned sensory selection may also contribute to the dynamic reshaping of sensory representations during attention.

Lateral propagation as a gain mechanism also has implications for downstream readout. If behavioral relevance is encoded partly through the spatial extent of cortical recruitment, then neurons outside the center response field ought to contribute to downstream decision-making even when they are not strongly tuned to the stimulus. Two recent studies provide causal evidence consistent with this idea (Bounds and Adesnik, 2025; Gauld et al., 2024). Using holographic photostimulation, both studies showed that recruiting sensory cortical neurons that were not strongly stimulus-encoding could bias decision-making, emphasizing the importance of population recruitment over the tuning of individual neurons. Lateral propagation could therefore provide an endogenous mechanism for regulating the number and spatial extent of cortical neurons recruited to influence downstream decision-related populations.

The precise circuit mechanisms underlying target-evoked propagation and distractor-evoked restriction remain to be determined. A notable finding of this study is that wMC actively suppresses distractor-evoked wS1 activation and lateral propagation (Fig. 6). Suppression of wMC not only increased distractor-evoked activity within the center response field and across the broader cortical field, but also significantly altered the peak-spread relationship, indicating a disproportionate increase in broader cortical recruitment relative to center region activity. These observations provide a candidate neural substrate for our previous finding that suppression of wMC increases distractor detection (Zareian *et al*., 2023; Zhang and Zagha, 2023). We recently demonstrated that distractor_wMC preferentially activates GABAergic interneurons within distractor_wS1 (Zhang and Zagha, 2023), consistent with a net inhibitory top-down influence. Additional experiments will be required to identify the cell-types mediating this suppression and to determine whether limiting lateral propagation is necessary for distractor filtering.

Our simple two-dimensional computational model tested whether modulation of lateral interactions is sufficient to reproduce key features of target-selective sensory processing. Increasing lateral gain reproduced several target-distractor differences, including increased prestimulus activity, mildly increased center-evoked activity, robustly increased full-field activity, and an altered peak-spread relationship. In contrast, within this model, alternative gain manipulations did not reproduce the observed relationship between response magnitude and spatial spread. These findings do not exclude other gain mechanisms, but demonstrate that modulating lateral interactions is sufficient to reproduce multiple spatial response features. Notably, the lateral gain model did not reproduce the sustained increase in target-evoked wS1 activity (Supplemental Fig. 1), indicating that additional mechanisms likely contribute to target-distractor differentiation.

Together, these findings expand the concept of sensory gain from modulation of responses within a stimulus representation to modulation of the cortical territory recruited by that representation. By regulating lateral propagation, cortical circuits may control the neuronal population engaged to influence downstream processing. Such regulation provides a complementary mechanism for selectively expanding the cortical territory recruited by behaviorally relevant stimuli while restricting that recruited by distractors.

## Materials and Methods

### Datasets

This study combines new experimental datasets, new analyses of previously acquired datasets, and computational modeling. *Previously acquired datasets:* Single-unit recordings from target-aligned and distractor-aligned whisker primary somatosensory cortex (target_wS1 and distractor_wS1; Supplemental Figs. 1 and 2) were acquired as part of a previously published study (Zhang and Zagha, 2023). The analyses of mean firing rate and stimulus encoding shown in Supplemental Fig. 1 extend analyses of these previously acquired data, whereas the Spike Time Tiling Coefficient (STTC) analyses in Supplemental Fig. 2 were performed specifically for the present study. Widefield Ca²⁺ imaging datasets used for the expert-session spatial analyses (Fig. 2) and longitudinal learning analyses (Fig. 3) were also previously acquired (Aruljothi *et al*., 2020; Marrero *et al*., 2024; Marrero *et al*., 2021), but the radial-profile, peak–spread, fractional-activation, and related spatial analyses reported here are new. *New datasets:* The laminar electrophysiological recordings and current-source-density analyses in Fig. 4 and Supplemental Fig. 4, the electrophysiological and video-based temporal-alignment analyses in Fig. 5 and Supplemental Fig. 5, and the widefield imaging experiments performed during pharmacological suppression of distractor-aligned whisker motor cortex (distractor_wMC) in Fig. 6 and Supplemental Fig. 6 were newly acquired for this study. The numerical simulations in Supplemental Fig. 3 and recurrent-network simulations in Fig. 7 and Supplemental Fig. 7 were also newly performed for the present study.

### Animal subjects

All procedures were approved by the Institutional Animal Care and Use Committee at the University of California, Riverside and were performed in accordance with institutional and NIH guidelines. Male and female adult mice were used. Electrophysiological experiments used C57BL/6J mice (Jackson Laboratory, #000664) and, for a subset of experiments, VGAT-ChR2 mice (Jackson Laboratory, #014548); the optogenetic transgene was not exploited in the experiments reported here. Widefield Ca²⁺ imaging experiments were performed in mice expressing GCaMP6s pan-neuronally (Snap25-2A-GCaMP6s-D, Jackson Laboratory, # 025111). Mice were maintained on a 12 h light/dark cycle with unrestricted access to food. During behavioral training, water access was controlled and body weight was maintained at ≥85% of free-feeding weight.

### Surgical procedures and head fixation

Surgical procedures for head fixation and chronic imaging preparation were performed as previously described (Aruljothi *et al*., 2020; Marrero *et al*., 2024; Zareian *et al*., 2023; Zareian *et al*., 2021; Zhang and Zagha, 2023). Briefly, anesthesia was induced with ketamine (100 mg/kg) and xylazine (10 mg/kg, intraperitoneal) and maintained as necessary with 1–2% isoflurane. The dorsal skull was exposed and a stainless-steel headpost containing an approximately 8 × 8 mm central imaging window was attached using cyanoacrylate adhesive. For widefield imaging, the exposed skull within the window was covered with a transparent silicone elastomer to permit repeated visualization of dorsal cortex. Meloxicam (0.3 mg/kg) and enrofloxacin (5 mg/kg) were administered postoperatively according to the previously described protocol. Animals recovered for at least 3 days before water restriction and behavioral training. For acute laminar electrophysiological recordings, a small craniotomy and durotomy were made over wS1 or at defined positions medial to wS1 immediately before electrode insertion. For pharmacological suppression experiments, an additional small craniotomy was made over distractor-aligned whisker motor cortex for local muscimol delivery as described below.

### Selective whisker-detection task

Mice were trained in a head-fixed whisker-based Go/NoGo selective-detection task previously described in detail (Aruljothi *et al*., 2020; Marrero *et al*., 2024; Marrero *et al*., 2021; Zareian *et al*., 2023; Zareian *et al*., 2021; Zhang and Zagha, 2023). Behavioral control was implemented using custom MATLAB and Arduino routines. Whisker stimuli were delivered by piezoelectric actuators coupled to small paddles positioned within the D2–D4/E2–E4 whiskers on opposite sides of the face. The target and distractor stimuli were matched in amplitude and duration within each mouse and consisted of single rapid whisker deflections with a rising phase of 50–100 ms and paddle velocity of 10–20 mm/s.

Trials began following a variable 5–9 s intertrial interval. Target, distractor, and catch trials were interleaved. Stimulus onset was followed by a 200-ms response lockout period and then a 1-s response window. Licking during the intertrial interval or lockout period reset the intertrial interval, and trials containing licking prior to the response window were excluded from the behavioral and neural analyses described here. A lick during the response window following a target stimulus was classified as a hit and produced water reward; failure to lick was classified as a miss. Licking following a distractor stimulus was classified as a false alarm and produced a timeout, whereas withholding licking was classified as a correct rejection and was rewarded with a subsequent target stimulus. Following a hit, miss, or false alarm, the subsequent trial had a 20% chance of being a target stimulus. Water rewards were approximately 5 µL.

Target–distractor discrimination was quantified using signal-detection theory as:

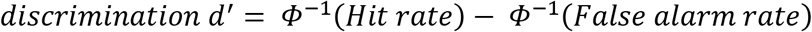

where Φ⁻¹ denotes the inverse cumulative normal distribution. To avoid infinite d′ values when response rates were 0 or 1, we applied a one-trial correction before the inverse-normal transformation, such that corrected response rates were calculated as (number of responses + 0.5)/(number of trials + 1). Animals were considered expert after achieving d′ > 1 on at least three consecutive training sessions. For stimulus-specific detection analyses relative to catch trials, detection was calculated independently for each whisker stimulus as:

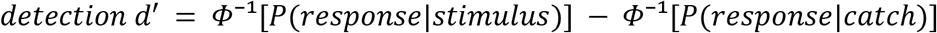

Unless otherwise indicated, neural analyses focused on the 200 ms lockout period between stimulus onset and onset of the response window, thereby reducing contamination by licking responses.

### Single-unit electrophysiology and stimulus encoding

Previously acquired wS1 single-unit recordings were analyzed as described in (Zhang and Zagha, 2023). Briefly, recordings were obtained with silicon probes containing tetrode-clustered recording sites (Neuronexus A1x16-Poly2-5mm-50s-177 or equivalent), sampled at 32 kHz. Signals used for spike detection were band-pass filtered at approximately 600–6000 Hz, and single units were identified by offline spike sorting according to previously described criteria. For the present study, analyses were restricted to neuronal populations recorded within the somatotopic wS1 representation of the presented whisker stimulus: target-aligned wS1 neurons during target trials (target_wS1) and distractor-aligned wS1 neurons during distractor trials (distractor_wS1). Neurometric d′ quantified the separability of prestimulus and post-stimulus spike-count distributions as previously described (Zhang and Zagha, 2023), yet performed here on single unit data instead of collapsed multiunit data.

### Spike synchrony analysis

Pairwise spike synchrony was quantified using the Spike Time Tiling Coefficient (STTC), which measures the temporal association between two spike trains while reducing the dependence of synchrony estimates on firing rate (Cutts and Eglen, 2014). For a pair of simultaneously recorded units A and B:

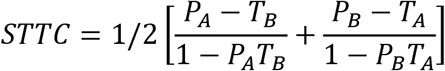

where *P_A_* and *P_B_* are the proportions of spikes in each train occurring within +/- Δt of a spike in the other train, and *T_A_* and *T_B_* are the fractions of the recording interval lying within +/- Δt of spikes in each respective train. Δt was tested at 5, 15, and 25 ms. STTC was calculated for simultaneously recorded unit pairs and averaged within recording sessions. Chance synchrony was estimated from 100 shuffled realizations generated by randomly permuting trial identity for one neuron relative to the other within each pair, thereby disrupting cross-neuron trial alignment while preserving within-trial spike timing; the 95% interval of the shuffled distribution was used for visualization. Session-level target_wS1 and distractor_wS1 STTC values were compared using two-sided unpaired t-tests. Within each population, real and shuffled session-level STTC values were compared using paired two-sided t-tests.

### Widefield Ca²⁺ imaging

Widefield Ca²⁺ imaging was performed as previously described (Aruljothi *et al*., 2020; Marrero *et al*., 2024). Two widefield imaging systems were used for data acquisition. Data presented in Figs. 2 and 3 were acquired using a system in which GCaMP6s fluorescence was excited with a 470-nm LED (Thorlabs M470L3), collimated through an ACL2520-A lens, and passed through a Chroma ET480/40x excitation filter. Excitation and emission light were separated by a T510lpxrxt dichroic mirror. Images were acquired through a Macroscope IIa with a 75-mm inverted lens at 0.7× magnification and recorded at 10 Hz using an RT sCMOS camera (SPOT Imaging). Data presented in Fig. 6 were acquired using a second system based on the protocol described by (Couto et al., 2021). GCaMP6s fluorescence was excited with the same 470-nm LED and passed through a Chroma ET470/40x excitation filter. Excitation and emission light were separated by a Chroma T495lpxr long-pass dichroic mirror. Images were acquired through a Nikon 85-mm objective lens and a Nikon 105-mm imaging lens and recorded at 10 Hz using a PCO Edge 5.5 sCMOS camera (PCO).

Imaging began 1 s before stimulus onset and continued through the behavioral response period for a total trial duration of approximately 2.2 s. For each trial and pixel, baseline fluorescence was calculated from the 1-s prestimulus period. ΔF/F₀ was calculated relative to this prestimulus baseline as described previously (Aruljothi *et al*., 2020). Neural stimulus-encoding maps were then generated independently for target and distractor trials by comparing prestimulus and post-stimulus fluorescence distributions at each pixel using receiver-operating-characteristic analysis. ROC area was transformed to neurometric d′ as

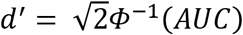

Unless otherwise indicated, spatial analyses used the second post-stimulus imaging frame, during which robust stimulus-related activation was consistently observed, and which fell entirely within the response lockout period.

For across-session visualization, imaging fields were registered using bregma and the bilateral wS1 representations as anatomical/functional landmarks, consistent with the Allen Mouse Common Coordinate Framework (Wang et al., 2020). Quantitative analyses were performed on individual session maps before cross-session averaging.

### Spatial and radial analysis of cortical activity

Spatial propagation was quantified from session-level neurometric d′ maps using custom MATLAB routines. Analyses were restricted to the parieto-occipital imaging field caudal to bregma, thereby excluding frontal cortex. For each target or distractor map, the maximum finite d′ value was identified within the expected somatotopic wS1 region for that stimulus. This position defined the center of the radial analysis.

For the expert imaging dataset (Fig. 2), the effective pixel size was 41 µm and concentric radial regions were constructed in 5-pixel increments, corresponding to 205 µm per radial bin.

The first radial region, 0 < r ≤ 205 µm, was designated the center region. Mean activity was calculated independently for each annulus from all finite pixels within that distance range. The center-three-regions measurement combined the first three radial regions using a pixel-weighted mean. Radial profiles were averaged across sessions. For normalized radial profiles, the group-averaged radial profile for each condition was divided by that condition’s mean center region value. Center region activity was the mean neurometric d′ within the first radial region; ipsilateral activity was the mean across the portion of the parieto-occipital field containing the relevant wS1 center; full field activity was the mean across all finite pixels in the bilateral parieto-occipital analysis field.

To quantify the relationship between local response magnitude and broader cortical recruitment, center region activity was plotted against full field activity for each recording session (which we refer to as the peak-spread relationship). Ordinary least-squares regression was used to describe each condition separately. To test across conditions, a linear mixed-effects model was fit with full field activity as the dependent variable, centered center region activity, condition, and their interaction as fixed effects, and session as a random intercept:

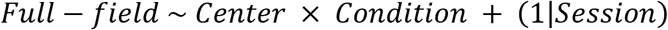

The Center × Condition interaction tests whether the peak-spread slopes differ between target and distractor trials.

### Gaussian simulations of peak–spread relationships

To illustrate the distinction between multiplicative response scaling and changes in spatial spread (Supplemental Fig. 3), idealized Gaussian activity profiles were generated numerically. In the multiplicative-gain simulation, Gaussian amplitude was scaled over a range of values while spatial width was held constant. In the spread-gain simulation, Gaussian width was increased while amplitude scaling was otherwise preserved. Peak activity and integrated spatial activity were then calculated for each simulated profile.

### Longitudinal analysis across task learning

Widefield imaging data acquired longitudinally during behavioral learning were reanalyzed for the present study (Fig. 3). Six mice contributed imaging sessions spanning naïve, intermediate, and expert performance. Sessions were categorized according to target-distractor behavioral d′ as naïve (d′ < 0.8), intermediate (0.8 < d′ < 1.2), or expert (d′ > 1.2). Neurometric maps and spatial metrics were generated using the same procedures described above. When multiple sessions from an individual mouse fell within a behavioral stage, session measurements were averaged within mouse and stage for mouse-level comparisons.

Center-region and full-field encoding were analyzed using two-factor repeated-measures ANOVA with trial type (target/distractor) and training stage (naïve/intermediate/expert) as within-subject factors. Learning-dependent changes in the peak-spread relationship were assessed with mixed-effects regression in which full field encoding was modeled as a function of center region encoding and training stage. For the combined target/distractor analysis, trial type and its interactions were additionally included; the Center × Trial type × Training stage interaction tested whether learning changed the peak-spread slope differently for target and distractor stimuli.

### Laminar electrophysiological recordings

New laminar recordings were acquired through 16-channel NeuroNexus electrode arrays in either single-shank linear probe (A1x16, 100 μm site spacing) or 4-shank polytrode configurations (A4x4, 200 μm site spacing). On recording days, expert mice were anesthetized under isoflurane for establishing and maintaining craniotomies and were allowed to recover from anesthesia for at least one hour within their home cage. Recovery was assessed based on activity levels in the home cage and regular behavioral performance during the first minutes of the task. When implanted, both electrode array configurations spanned superficial and deep cortical layers: A1x16, full cortical depth, A4x4, from 200-800 µm from pia.

For A1x16 probe implants, craniotomies and durotomies (∼0.5 mm diameter) were established above the wS1 center response field (from bregma: 3.0 mm lateral, 1.0-1.5 mm posterior) or the medial parietal region (from bregma: 1.5 mm lateral, 0.4-1.0 mm posterior). For A4x4 polytrode implants, craniotomies and durotomies spanned approximately 0.8 mm to accommodate the 200μm spacing between shanks (0.6 mm across all four shanks). The aforementioned wS1 coordinates were used for the A4x4 recordings, and the shanks were implanted such that the most lateral shank was placed 3.0 mm lateral of bregma. The remaining shanks were placed 2.8 mm, 2.6 mm, and 2.4 mm lateral of bregma.

Once recovery was established, probes were slowly guided into the brain using Narishige hydraulic micro-manipulators. For all wS1 recordings, whisker alignment was performed during probe implantation and verified by post-hoc analysis. Prior to recording, individual whiskers were mapped by hand while observing live LFP displays on the acquisition software. Post-hoc verification consisted of plotting stimulus-evoked LFP activity and confirming robust responses that peaked within 30ms after stimulus onset. Sessions that did not satisfy these criteria were excluded. Broadband signals were acquired through Neuralynx software at a rate of 32kHz and were recorded simultaneously with behavioral events. For analysis, LFP signals were low-pass filtered at 160Hz and downsampled to 200Hz.

Current-source density (CSD) was extracted from recordings acquired with A1x16 linear probe only. From LFP signals, CSD was estimated as the negative second derivative across adjacent recording channels:

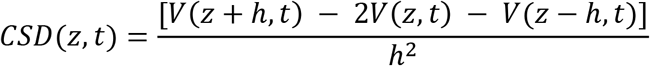

where *h* is the 100 µm interelectrode spacing.

Latencies plotted in the linear regression were sourced from all laminar recordings, regardless of probe used. LFP response onset latency was measured relative to stimulus onset from trial-averaged traces as the first of two consecutive samples that are three standard deviations above the average of the prestimulus window. In Fig. 4J, the position of two datapoints were slightly shifted for visualization purposes only. Propagation velocity was estimated by ordinary least-squares regression of LFP onset latency against medial distance from the wS1 center. Velocity was calculated as the inverse of the regression slope.

### Video motion-energy and reaction-time alignment analyses

Behavioral video was acquired synchronously with neural and task events at 55 frames/s. Rectangular regions of interest were manually positioned over the stimulus paddle, unstimulated whiskers of the target whisker field, and tongue/jaw region (Supplemental Fig. 5). Motion energy (ME) was calculated separately for each region of interest (ROI) as the sum of the absolute pixel-wise change in image intensity between consecutive video frames. The resulting ME trace was z-scored independently for each ROI across the full recording session before trial parsing and alignment to behavioral events. Temporal-alignment analyses were performed on Hit trials. Within each recording session, Hit trials were divided into fast, medium, and slow reaction time terciles of that session’s reaction-time distribution. Neural and motion-energy signals were then averaged separately for each tercile. The same underlying trials were analyzed either relative to stimulus onset or after realignment to the behavioral response. Latencies were determined as the time to half-maximum activation; for Fig. 5F, the latencies of the fast reaction time terciles were used. Fast-tercile response latencies were compared across signal types using a linear mixed-effects model with signal type as a fixed effect and dataset as a random intercept. The overall effect of signal type was assessed using a marginal F-test, followed by Holm-corrected pairwise comparisons.

To quantify whether each signal was more tightly aligned to stimulus onset or to the behavioral response, we first calculated the standard deviation of the fast-, medium-, and slow-reaction-time tercile latencies under stimulus alignment and response alignment for each dataset. We then calculated a paired-samples Cohen’s *d_z_* as:

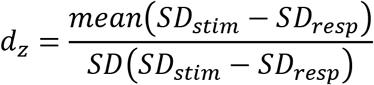

Positive values indicate tighter response alignment, whereas negative values indicate tighter stimulus alignment.

### Pharmacological suppression of distractor-aligned whisker motor cortex

To determine whether lateral sensory recruitment is regulated by frontal cortex, widefield Ca²⁺ imaging was performed before, during, and after local pharmacological suppression of distractor-aligned whisker motor cortex (distractor_wMC) in expert mice (Fig. 6 and Supplemental Fig. 6). distractor_wMC was identified at approximately (from bregma: 1 mm anterior, 1 mm lateral). Muscimol (2 mM, 250 nL) was delivered at a depth of 700 µm through a glass micropipette using a Nanoject III Programmable Nanoliter Injector (Drummond Scientific) at a rate of 1 nL/s. Imaging during suppression began approximately 30 min after injection. Baseline recordings were obtained 1 day before muscimol administration and recovery recordings were obtained 1-2 days after muscimol administration. For sham-control sessions (baseline and/or recovery), the same surgical procedures were applied but without muscimol injection. Specifically, mice were maintained under isoflurane anesthesia for approximately 45 min and a craniotomy was performed over distractor_wMC if necessary. Behavioral testing and imaging began 30 min after the sham manipulation. Target and distractor neurometric d′ maps were calculated using the same procedures described above. Quantification was restricted to cortex caudal to bregma. Differences in center-region and full-field encoding and behavioral detection were tested with paired two-sided t-tests. Differences in peak-spread slopes were tested separately for target and distractor stimuli using linear mixed-effects models with full-field activity as the dependent variable, center-region activity, drug condition, and their interaction as fixed effects, and mouse as a random intercept:

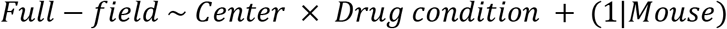

where the Center × Drug-condition interaction tested whether muscimol altered the relationship between center activity and broader cortical recruitment. Significance of fixed effects was assessed using marginal F tests with Satterthwaite degrees of freedom.

For visualization of the spatial extent of distractor_wMC suppression (Supplemental Fig. 6), muscimol minus pre-muscimol difference maps were calculated during the third post-stimulus imaging frame. Negative values indicate reduced activity during muscimol application.

### Recurrent-network simulations

We simulated a two-dimensional recurrent rate network consisting of a 41 x 41 lattice of units, although analyses focused on the center 11 x 11 units. The complete network dynamics followed:

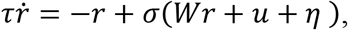

where *r*(*t*) is an *N* × 1 vector of all neurons’ rates at time *t* (*N* = 41^2^ = 1681 is the number of neurons), *τ* = 0.1 is the network time constant, *W* is an *N* × *N* sparse matrix of recurrent weights, *u*(*t*) = *u*_bg_ + *u*_stim_(*t*) is the vector of total (background plus stimulus-evoked) exogenous inputs to each neuron, *u*_bg_ = 1 is a uniform tonic input for all neurons, *u*_stim_(*t*) = *A* only for the center neuron during stimulus presentation and *u*_stim_(*t*) = 0 otherwise, *η*(*t*) ∼ *N*(0,0.01) is a vector of independent Gaussian noises perturbing all neurons, and *σ*(*z*) = max(*z* − *θ*, 0)*^p^* is a thresholded and rectified supra-linear activation function with *θ* = 0.7 and *p* = 1.5. Each unit interacted with itself and with neighboring units defined by Chebyshev distance. Baseline self-coupling was *W_s_* = −1, coupling to units at distance 1 was *W_d_*_1_ = 0.10, and coupling to units at distance 2 was *W_d_*_2_ = 0.05. Dynamics were integrated with a fixed-step Euler integration with *dt* = 0.01 for *T* = 15 s. The focal sensory input was applied to the center unit beginning at *t* = 5s and lasted 200 ms under the baseline condition. Five sensory-input amplitudes of *A* = 1,2, … 5 a.u. were used to generate peak-spread relationships.

Four manipulations were compared with the baseline network. Center gain was implemented by increasing only the self-coupling of the center unit from *W_s_* = −1 to 0. Lateral gain was implemented by selectively increasing excitatory coupling to and from the center unit, with center-associated distance-1 connections increased from *W_d_*_1_ = 0.10 to 0.20 and distance-2 connections from *W_d_*_2_ = 0.05 to 0.10. An increased stimulus condition increased external sensory-input amplitude by 2x, without changing recurrent connectivity. An increased duration condition increased stimulus duration by 2x while preserving its amplitude and network connectivity.

Prestimulus activity was calculated from the 200 ms interval immediately preceding stimulus onset. Stimulus-evoked activity was calculated by subtracting the mean prestimulus activity from the mean activity during the post-stimulus analysis window (0.1-0.3 s after stimulus onset), separately for each lattice node. Peak-spread relationships were generated by regressing spatial recruitment against center activity across the five stimulus amplitudes. The model comparisons were intended to test qualitative mechanistic predictions rather than to fit free parameters to the experimental data.

### Quantification and statistical analysis

Analyses were performed using custom MATLAB scripts (MathWorks). All statistical tests were two-sided, and α = 0.05 was used as the criterion for statistical significance. Data are reported as mean +/- SEM unless otherwise indicated. Sample sizes, statistical tests, test statistics, and p-values are reported in the Results.

### Data and code availability

The data supporting the findings of this study and the custom analysis and simulation code used to generate the principal results will be made publicly available upon publication.

## Acknowledgements

This work was supported by National Institutes of Health (NIH) grants R01NS148491 (E.N. and E.Z.) and R01NS112200 (H.Y. and E.Z.).

---

**Supplemental Figure 1.**
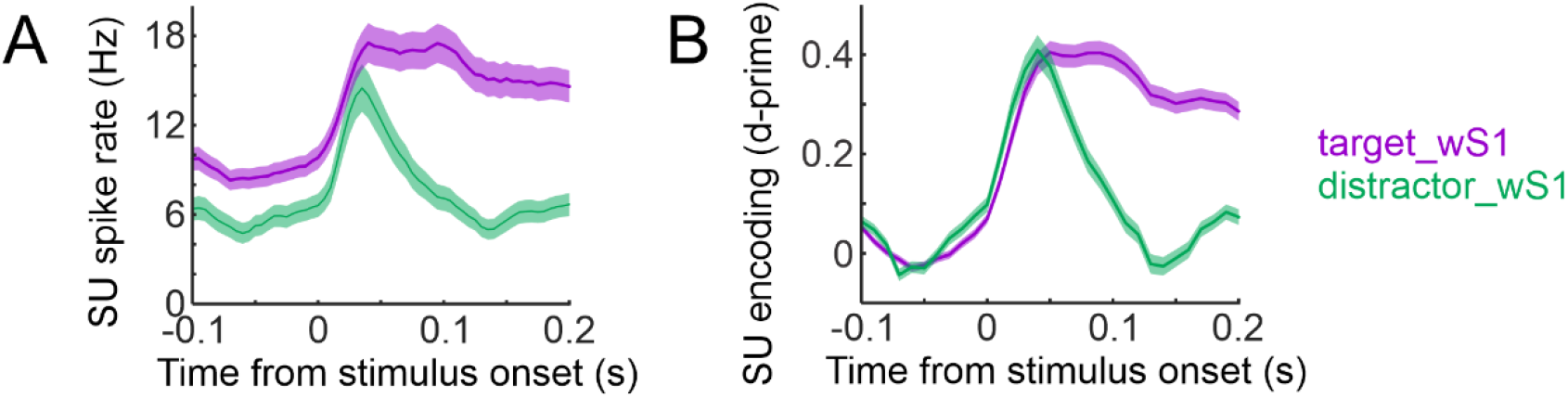
Target and distractor stimuli evoke similar peak sensory encoding within their center response fields. (A) Mean single unit (SU) firing rates aligned to stimulus onset within the center response fields: target-aligned wS1 on target trials (target_wS1, purple) and distractor-aligned wS1 on distractor trials (distractor_wS1, green). (B) Neurometric stimulus encoding (d′) calculated from the same neuronal populations. Target and distractor populations exhibited similar rising phases and peak stimulus encoding. And yet, notable differences include higher prestimulus firing rates and sustained post-stimulus firing for the target compared to distractor population. Shading denotes SEM.

**Supplemental Figure 2.**
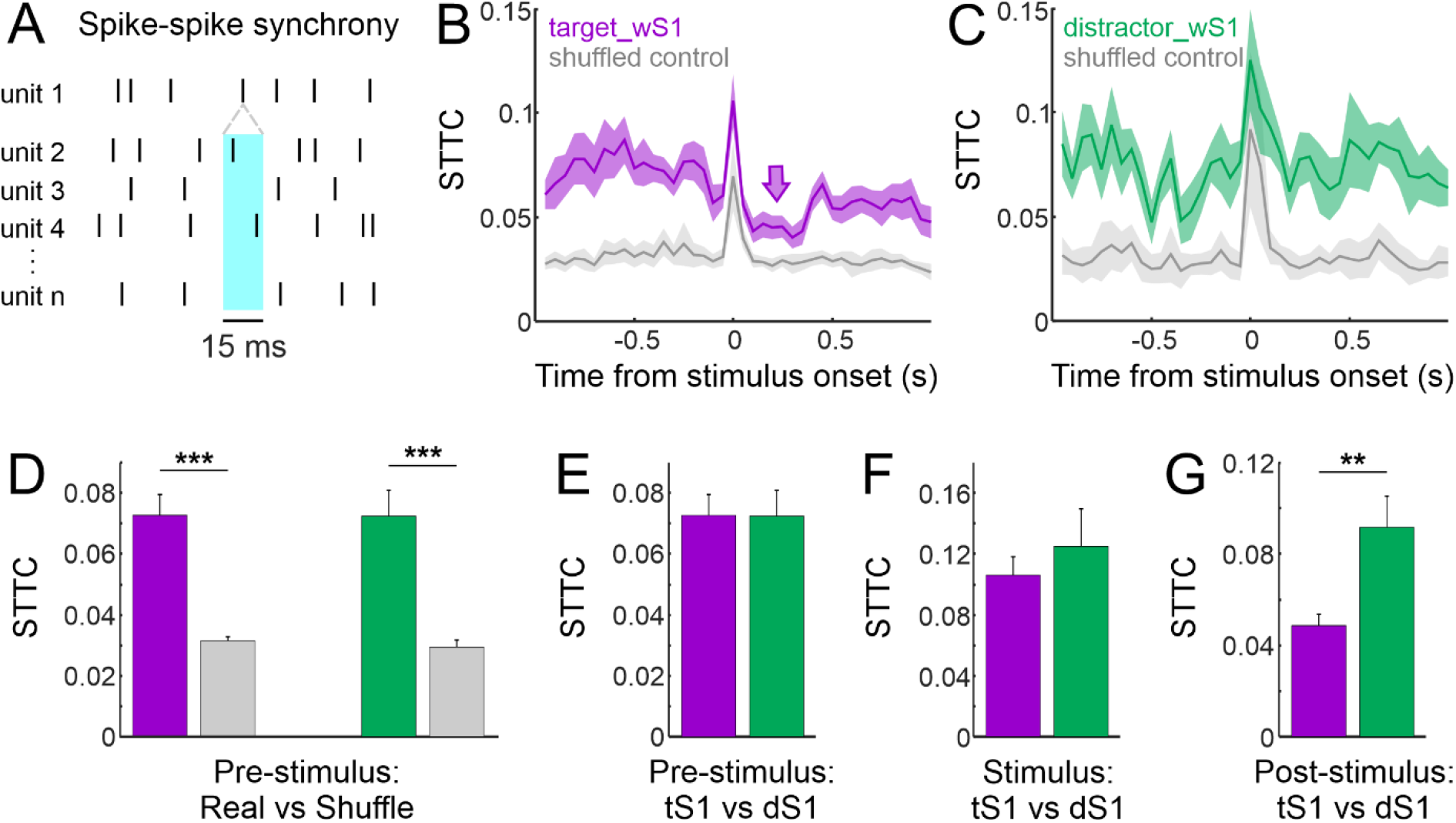
Spike synchrony is not selectively enhanced for target stimuli in sensory cortex. (A) Illustration of spike synchrony, as the co-occurrence of spikes from pairs of neurons within a defined time window. (B,C) Spike Time Tiling Coefficient (STTC) results averaged across sessions, for the target_wS1 condition [B, purple], the distractor_wS1 condition [C, green], and their shuffled controls (gray). Both target_wS1 and distractor_wS1 exhibit similar prestimulus spike synchrony, which transiently increases at stimulus onset. Notably, target_wS1 displayed reduced spike synchrony post-stimulus (purple arrow in B). (D) STTC comparison of real data (color) and shuffled controls (gray) prestimulus (-1 second to stimulus onset). Both target_wS1 and distractor_wS1 exhibit spike synchrony significantly above chance levels. (E-G) STTC comparisons between target_wS1 and distractor_wS1. STTC distributions are not different prestimulus (E) or at stimulus onset (F, single window -50 ms to +50 ms). Post-stimulus (G, +50 ms to +200 ms), distractor_wS1 spike synchrony is greater than target_wS1 spike synchrony. **, p<0.01; ***, p<0.001. Error bars and shade are SEM, except for the shade of the shuffled controls which are 95% confidence intervals.

**Supplemental Figure 3.**
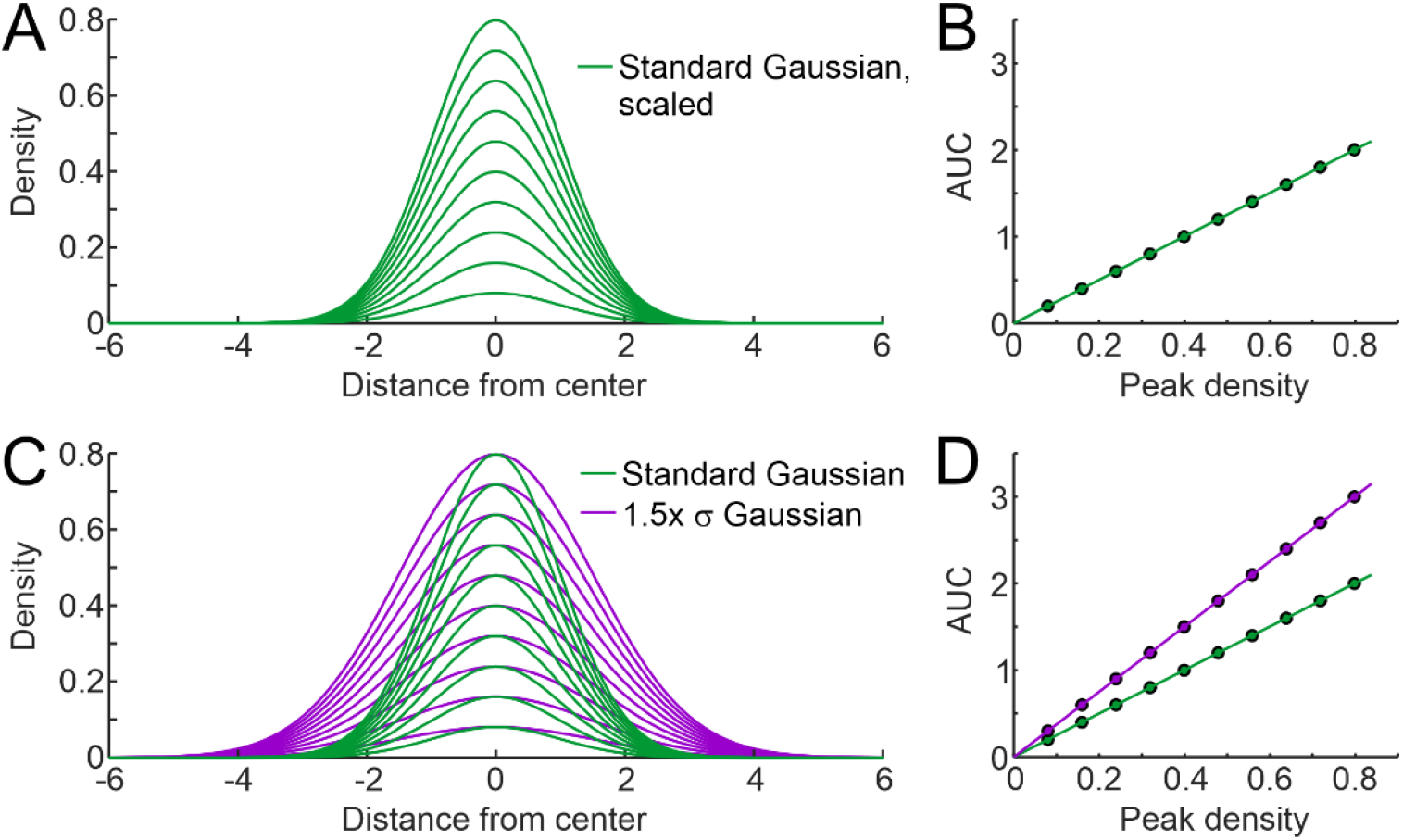
Lateral gain increases the spatial spread of activity relative to peak response. (A) Simulation of gain as multiplicative scaling. Gaussian spatial activity profiles generated across a range of scales. (B) Integrative activity (Area under the curve (AUC), y-axis) plotted against peak activity (x-axis) demonstrates a preserved peak-spread relationship across multiplicative scales. (C) Simulation of gain as increases in the width relative to the standard distribution (purple). (D) Increasing distribution width produces a steeper relationship between integrated and peak activity. These simulations illustrate the different predictions of multiplicative scaling versus modulations of spatial propagation.

**Supplemental Figure 4.**
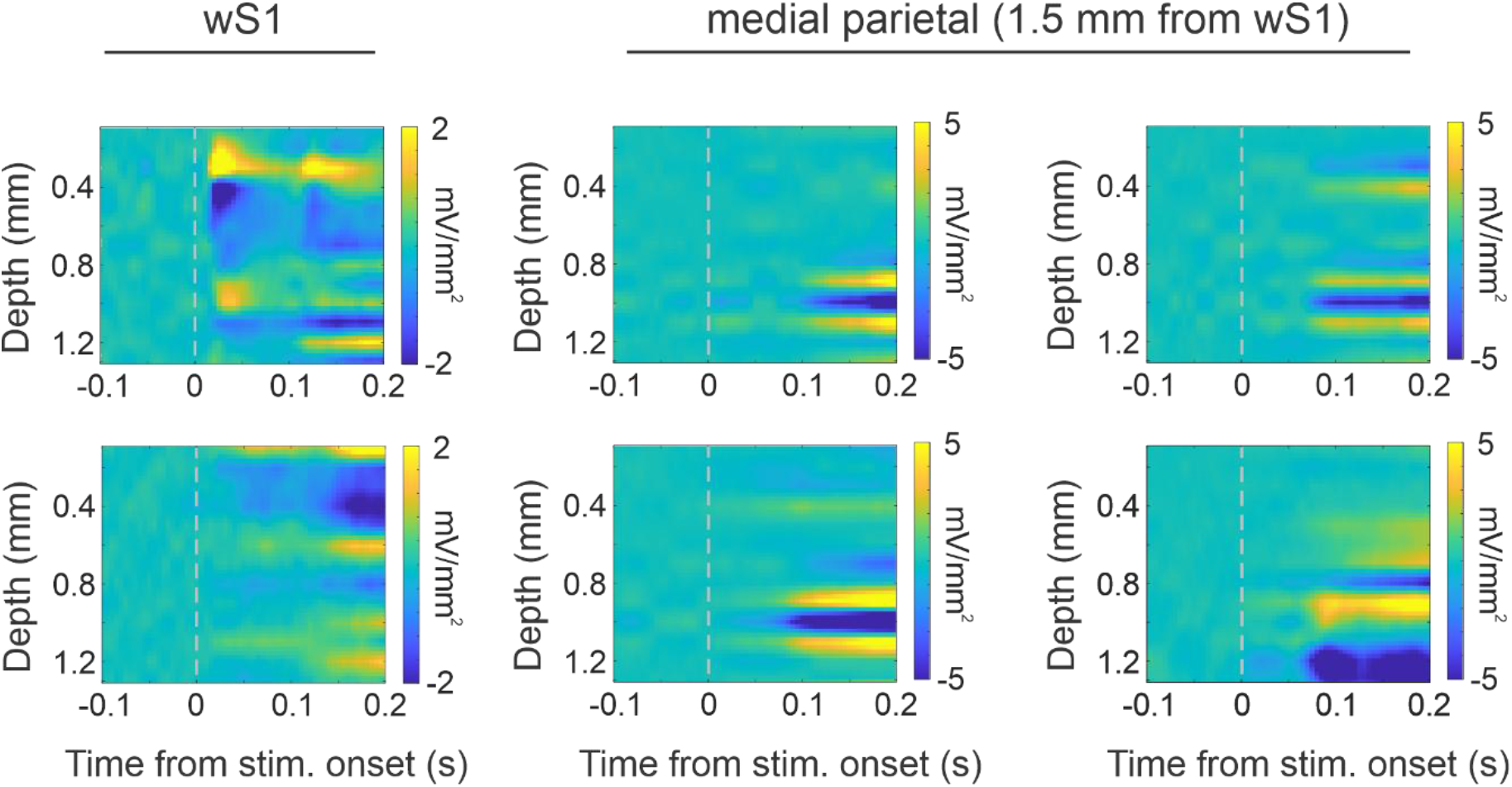
Laminar current-source-density profiles of target-evoked responses within wS1 and medial parietal cortex. Representative CSD maps from independent recording sessions within wS1 (left column) and 1.5 mm medial to wS1 (middle and right columns). Stimulus onset is indicated by the vertical dashed line. wS1 recordings show prominent stimulus-evoked current sinks at intermediate cortical depths, whereas recordings medial to wS1 show prominent sinks in deeper cortical layers, consistent with preferential recruitment of infragranular layers during lateral propagation.

**Supplemental Figure 5.**
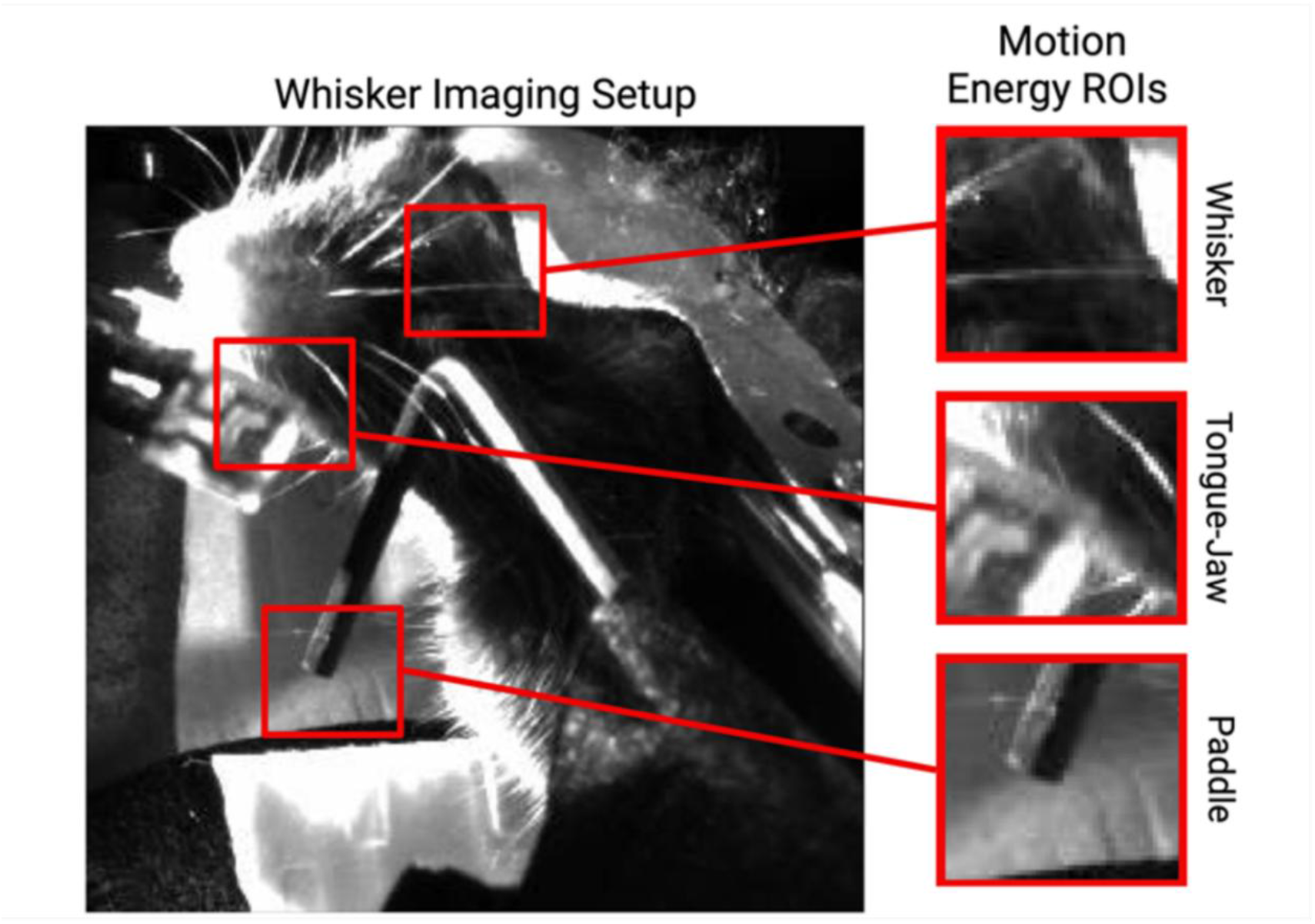
Video-based regions of interest used for motion energy analyses. Representative behavioral video frame showing the head-fixed mouse, stimulus paddle, and face. Red boxes indicate regions of interest used to quantify motion energy from the unstimulated whiskers, tongue/jaw region, and stimulus paddle. Enlarged examples of each region are shown at right.

**Supplemental Figure 6.**
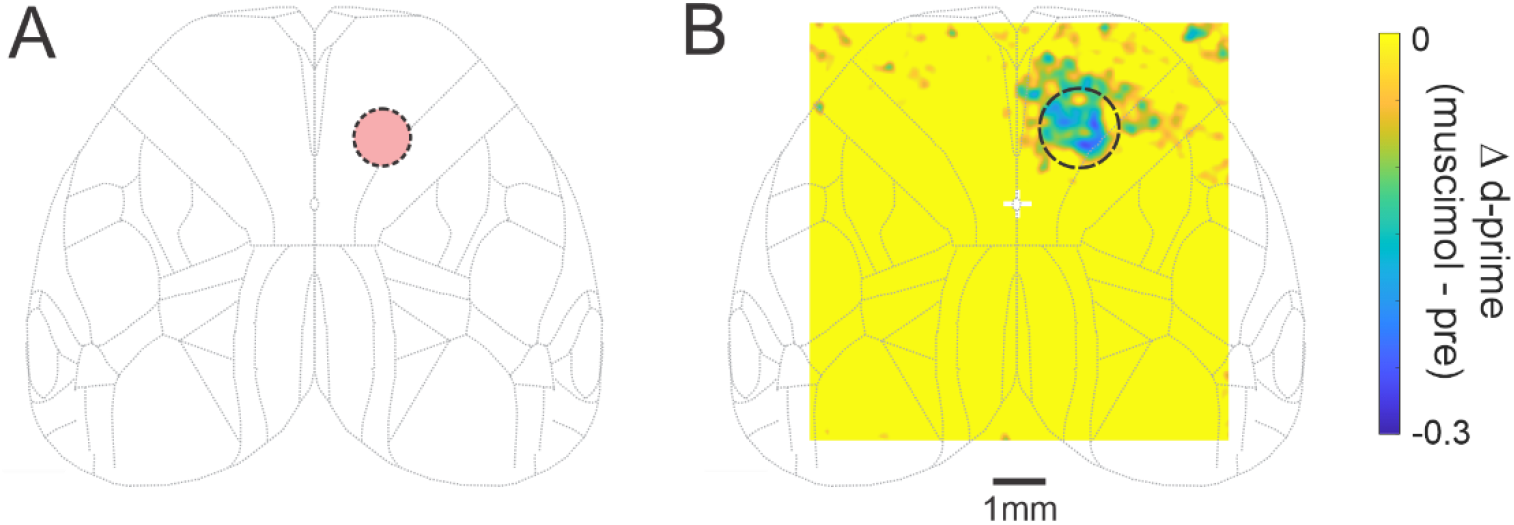
Location and spatial extent of distractor-aligned whisker motor cortex suppression. (A) Schematic showing the location of muscimol injection into distractor-aligned whisker motor cortex (distractor_wMC). (B) Grand average difference map of cortical activity during the early response window (third post-stimulus frame), calculated as muscimol minus pre-muscimol activity. Negative values indicate reduced activity during muscimol suppression. Dashed circle indicates the approximate distractor_wMC injection region.

**Supplemental Figure 7.**
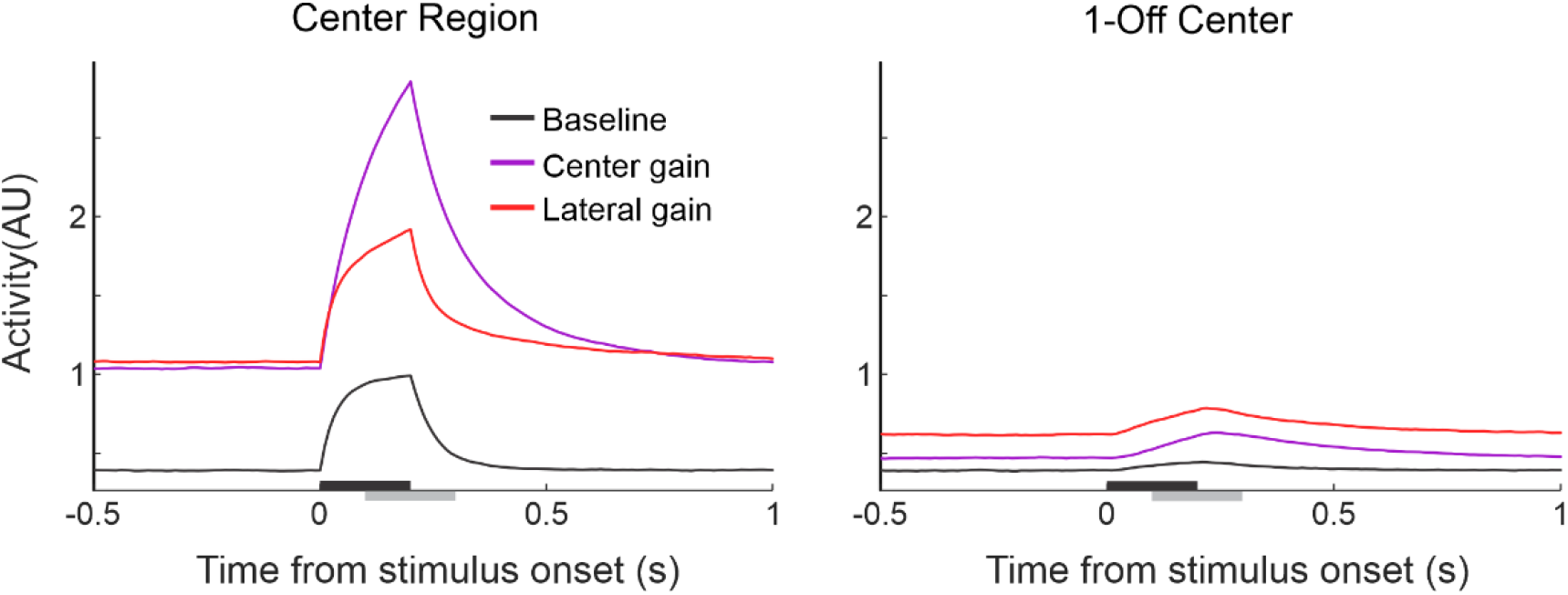
Center and lateral gain differentially affect center and surround activity. Simulated activity over time within the center region (left) and an off-center region (right) under baseline, center gain, and lateral gain conditions. Increasing center gain strongly amplifies activity at the stimulated center while producing comparatively modest changes in the surrounding network. Increasing lateral coupling produces a disproportionate enhancement of off-center activity, illustrating how lateral gain can alter the spatial distribution of stimulus-evoked activity rather than simply scaling the center response. The black bar overlying the x-axis denotes the time of stimulus presentation; the gray bar denotes the post-stimulus analysis window.

